# Rapid and synchronous chemical induction of replicative-like senescence via a small molecule inhibitor

**DOI:** 10.1101/2023.09.07.556710

**Authors:** Spiros Palikyras, Konstantinos Sofiadis, Athanasia Stavropoulou, Adi Danieli-Mackay, Vassiliki Varamogianni-Mamatsi, David Hörl, Simona Nasiscionyte, Yajie Zhu, Natasa Josipovic, Antonis Papadakis, Anne Zirkel, Aoife O’Connell, Gary Loughran, James Keane, Audrey Michel, Wolfgang Wagner, Andreas Beyer, Hartmann Harz, Heinrich Leonhardt, Grazvydas Lukinavicius, Christoforos Nikolaou, Argyris Papantonis

**Affiliations:** Institute of Pathology, University Medical Center Göttingen, Germany; Institute for Bioinnovation, Biomedical Sciences Research Center “Alexander Fleming”, Vari, Greece; Clinical Research Unit 5002, University Medical Center Göttingen, Germany; Faculty of Biology, Ludwig Maximilians University Munich, Germany; Cluster of Excellence on Cellular Stress Responses in Aging-associated Diseases (CECAD), University of Cologne, Germany; Center for Molecular Medicine Cologne, University and University Hospital of Cologne, Germany; EIRNA Bio (formerly Ribomaps), Cork, Ireland; Helmholtz-Institute for Biomedical Engineering, RWTH Aachen University Medical School, Germany; Institute for Stem Cell Biology, RWTH Aachen University Medical School, Germany; Department of NanoBiophotonics, Max Planck Institute for Multidisciplinary Sciences, Göttingen, Germany

**Keywords:** cellular ageing, chromatin, SASP, single cell genomics, 3D genome organization

## Abstract

Cellular senescence is now acknowledged as a key contributor to organismal ageing and late-life disease. Although popular, the study of senescence *in vitro* can be complicated by the prolonged and asynchronous timing of cells committing to it and its paracrine effects. To address these issues, we repurposed the small molecule inhibitor inflachromene (ICM) to induce senescence to human primary cells. Within six days of treatment with ICM, senescence hallmarks, including the nuclear eviction of HMGB1 and -B2, are uniformly induced across IMR90 cell populations. By generating and comparing various high throughput datasets from ICM-induced and replicative senescence, we uncovered significant similarity of the two states. Notably though, ICM suppresses the proinflammatory secretome associated with senescence, thus alleviating most paracrine effects. In summary, ICM induces a senescence-like phenotype rapidly and synchronously thereby allowing the study of its core regulatory program without any confounding heterogeneity.

## Introduction

From the onset of development until late-life stages, human cells encounter multiple signalling and stress cues. Many of these can lead to the induction of senescent phenotypes that commit cells to an irreversible growth arrest and are inextricably linked with ageing^1–3^. In fact, clearing senescent cells *in vivo* leads to prolonged health-and lifespan^4–7^. Depending on the initial trigger, senescent responses can be grouped into three types: replicative senescence (RS) occurring via telomere attrition, oncogene induced-senescence (OIS) due to oncogenic activation or senescence induced by genotoxic stress (e.g., irradiation)^8^. All give rise to distinct gene expression programs, which however converge to an underlying transcriptional signature linked to cell cycle regulation and transcriptional remodeling^9^.

Apart from the pronounced cell cycle arrest, there are multiple genomic hallmarks of the commitment to senescence. For example, models of OIS show formation of large senescence-associated heterochromatic foci (SAHFs)^10^, which involve the dissociation of heterochromatin from the lamina, the redistribution of Lamin B1^11,12^ and nuclear pore components^13^, as well as the interplay between DNMT1 and HMGA2^14^. These effects are also reflected on changes in the three-dimensional (3D) organization of chromosomes^14,15^, and many have also been recorded in a model for DNA damage-induced senescence^16^.

In RS, DNMT1 is linked to focal hypomethylation^17^, and HMGB (rather than HMGA) proteins seem to play a central role as they are quantitatively depleted from senescent cell nuclei^18^. The loss of HMGB1 was shown to affect both chromatin reorganization and mRNA splicing upon RS entry^19^, while that of HMGB2 was causal for heterochromatin imbalance and the formation of large senescence-induced CTCF clusters (SICCs)^20^. Cells maintained in RS long-term (i.e., “deep” senescence) display even more pronounced changes in 3D genome organization, mostly compaction of chromosomal arms and changes between compartments of active and inactive chromatin^21^ to suppress gene expression and activate transposable elements^22^. This is in line with spurious^23^ and accelerated transcription in senescence^24^, with an overall compromised ability to transcribe^20^, as well as with a transcription-dependent reorganization of chromatin loops^25^.

A key outcome of the senescent gene expression program is the production and secretion of a complex and cell type-specific mixture of proinflammatory factors: the senescence-associated secretory phenotype (SASP)^26–29^. SASP factors act in an autocrine and a paracrine manner^30^, and mediate both beneficial (e.g., wound healing) and detrimental effects of senescence (e.g., chronic inflammation and tumorigenesis)^31,32^. However, the production of SASP and other secondary signals (e.g., Notch in OIS^33^) by senescent cells emerging in a population can both promote and limit senescence spread^34^ in a manner that ultimately leads to a large heterogeneity of individual cell states^20,33,35,36^.

This pronounced heterogeneity, together with the asynchrony in senescence commitment by individual cells and the extended culture periods needed to reach replicative senescence complicate studies on the core of the senescent program that still remains only partially understood. Here, we address these caveats by the introduction of a novel and robust model of chemically-induced senescence via the repurposing of the small molecule inhibitor inflachromene (ICM)^37^. We show that ICM can induce senescence rapidly (within <6 days) and homogeneously in the popular fetal lung fibroblast (IMR90) cell model, while also constraining SASP production and its paracrine effects. We provide a comprehensive data resource by characterizing ICM-induced senescence to facilitate its adoption by the broader community.

## Results

### ICM induces a senescence-like phenotype in human fibroblasts

Inflachromene (ICM) was previously discovered as a direct binder of HMGB1/B2 proteins in a broad screen of compounds, and characterised as a potent blocker of their cytoplasmic translocation and extracellular release. As a result, ICM restricted inflammatory phenotypes *in vitro* and *in vivo* to exert a neuroprotective effect^37^. However, ICM was only tested in the neural context and only for up to 24 h. We subjected different isolates of donor-derived human lung fibroblasts (IMR90), one of the most popular models, for senescence studies^38–42^, to continuous exposure to different ICM concentrations. Treatment with 10 μM (but not 5 μM) ICM led to growth arrest within <4 days. Notably, removing ICM from the IMR90 growth medium after 6 days of treatment did not result in regrowth, while removal after 3 days of treatment did (Fig 1a). Such an effect by longer-term ICM treatment was unexpected and prompted us to ask whether it actually induced a senescent-like state to the cells.

**Fig 1.**
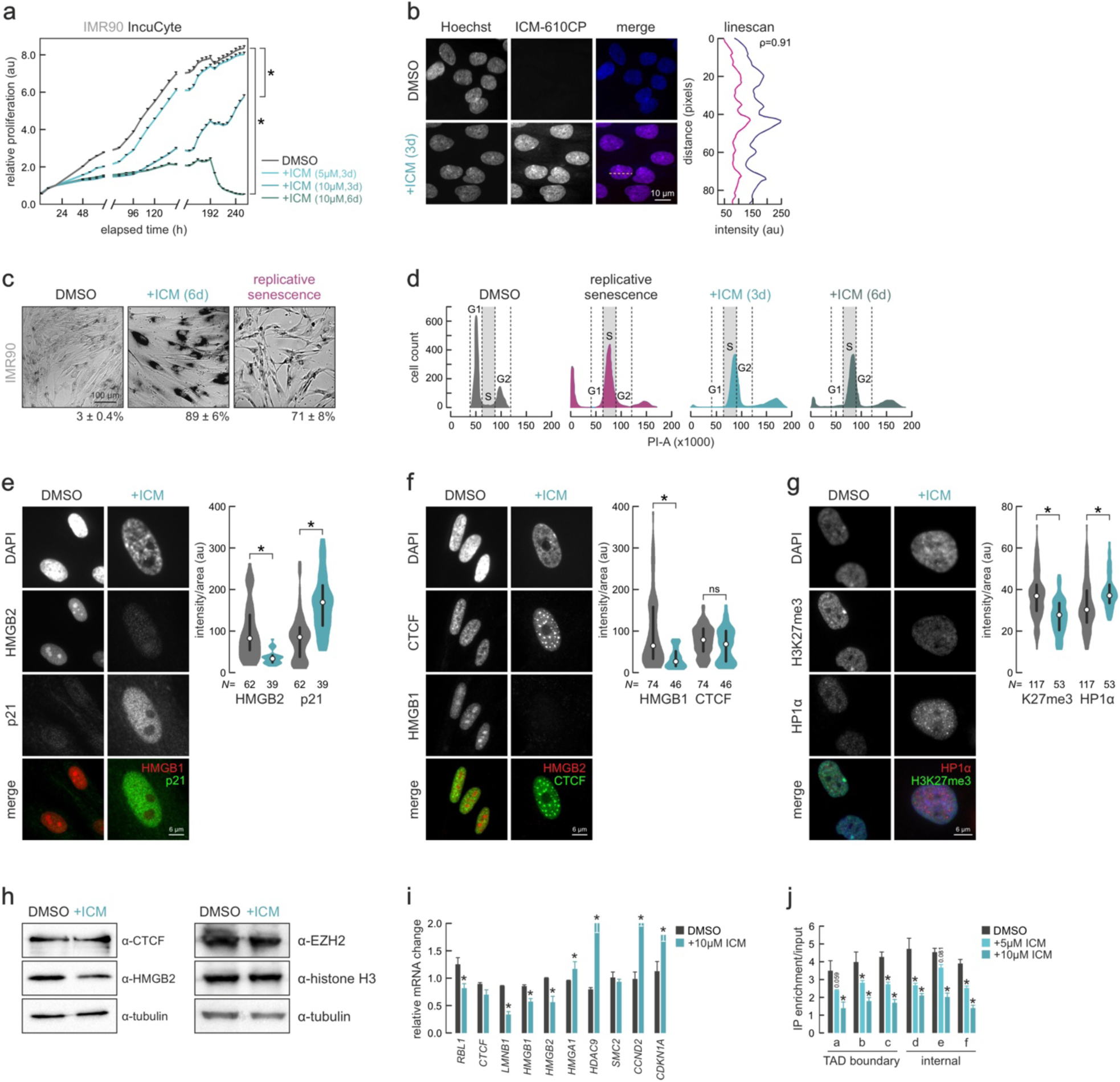
ICM treatment induces a senescent-like phenotype. **a.** Mean proliferation rates (±SEM from three independent replicates) of DMSO-and ICM-treated IMR90 for 3-6 days using automated live-cell imaging. \**P*<0.01, unpaired two-tailed Student’s t-test at 240 h. **b.** Representative widefield images of proliferating (top) and 610CP-C6-ICM-treated IMR90 (bottom) with DNA counterstained with Hoechst. The overlap between the two signals was assessed via a linescan. Bar, 5 μm. **c.** Proliferating, ICM-treated and senescent IMR90 assayed for β-galactosidase activity. ICM-treated and senescent cells appeared darker, indicative of their senescent state. **d.** FACS cell cycle profiles of PI-stained proliferating (DMSO), senescent or ICM-treated IMR90 for 3 and 6 days. **e.** Representative images of IMR90 showing treated or not with ICM for 6 days and immunostained for HMGB2 and p21. Bars, 6 μm. Violin plots (right) quantify changes in the levels of the two markers. *N*: number of cells analyzed per each condition. \**P*<0.05, two-tailed Wilcoxon-Mann-Whitney test. **f.** As in panel E, but immunostained for HMGB1 and CTCF. **g.** As in panel E, but immunostained for HP1α and H3K27me3. **h.** Western blot analysis of CTCF, HMGB2, EZH2 and histone H3 in proliferating (DMSO) and 6-day ICM-treated IMR90; α-tubulin levels provide a loading control Mean mRNA levels (±SD from two independent isolates) of selected senescence marker genes in proliferating (DMSO) and 6-day ICM-treated IMR90. \**P*<0.05, unpaired two-tailed Student’s t-test. **j.** Mean ChIP-qPCR enrichment levels (±SD from two independent isolates) at selected genomic positions (a-f) in proliferating (DMSO) and 3-or 6-day ICM-treated IMR90. \**P*<0.05, unpaired two-tailed Student’s t-test.

First, we synthesized a 610CP-C6-tagged version of ICM to verify that it indeed enters IMR90 nuclei and localizes to chromatin, where HMGBs reside. Both effects could be confirmed microscopically (Fig 1b). Next, we stained control and 6-day ICM-treated cells for β-galactosidase activity, a marker of senescence^43^. ICM-treated cells stained essentially uniformely SA-β-GAL-positive, even more than IMR90 from the same isolate driven to senescence by serial passaging (Fig 1c). FACS analysis showed that ICM treatment, already after 3 days, arrrested IMR90 in the late S-phase (24.4% and 29.2% after 3-and 6-day treatment, respectively, compared to 2.4% in DMSO-treated cells) comparably to the S-phase accumulation seen in replicatively senescent cells (23.6%; Fig 1d).

It has been established that the nuclear depletion of HMGB1 and HMGB2 from cell nuclei is a robust indicator of replicative senescence entry by different primary human cell types^19,20^. We could show that such pronounced nuclear loss of HMGBs is also achieved by ICM treatment of IMR90 (Fig 1e,f), together with the expected reduction in H3K27me3 levels (Fig 1g). These effects were coupled to the upregulation of senescence marker p21 (Fig 1e), the formation of senescence-induced CTCF clusters (SICCs; Fig 1f), and the emergence of HP1α foci (Fig 1g), again, much like what has been recorded for RS ^20^. Interestingly, all these changes could be detected, albeit to a somewhat smaller extent, upon treatment with ICM for 3 days (Supplementary Fig 1a), when growth arrest is not yet irreversibly committed to (Fig 1a). This suggests that HMGB1/-B2 loss and SICC formation are early events on the path to senescence, as previously postulated^20^.

These changes were also reflected in changes at the protein and/or mRNA levels of the factors involved, as well as of other senescence-regulated genes like *HDAC9*, *CCND2*, and *HMGA1* (Fig 1h,i). Finally, using ChIP-qPCR we confirmed loss of HMGB2 chromatin binding upon ICM treatment from known positions in RS-IMR90^20^ at both the boundaries of topologically-associating domains (TADs) and at other non-boundary regions (Fig 1j). In summary, short-term treatment of IMR90 with ICM results in irreversible growth arrest as well as in phenotypic changes likeing those of RS.

### Automated imaging and classification of nuclear features changes in ICM-treated cells

Replicative senescence induces similar changes to the nuclear morphology of different primary human cell types, includinig size increase and a characteristic pattern of DAPI chromatin staining ^19,20^. On the basis of these observations, we reasoned that ICM-treated cells could be classified as regards their senescence state via imaging of nuclear features. To this end, we devised an automated imaging and classification workflow to process images of >11,000 IMR90 cells that were either proliferating (early-passage), senescent, or treated with ICM for 3-9 days. In brief, our workflow used fixed cells counterstained by SiR-DNA to visualize chromatin. Cells were identified and imaged via automated confocal imaging, while super-resolution mid-planes of individual nuclei were also captured using the platform’s STED mode (Fig 2a). To discard erroneous detections, STED images of nuclei were filtered using machine learning-based quality control achieving 95% precision in identifying “good” versus “bad quality” images (Fig 2a; see Methods for details).

**Fig 2.**
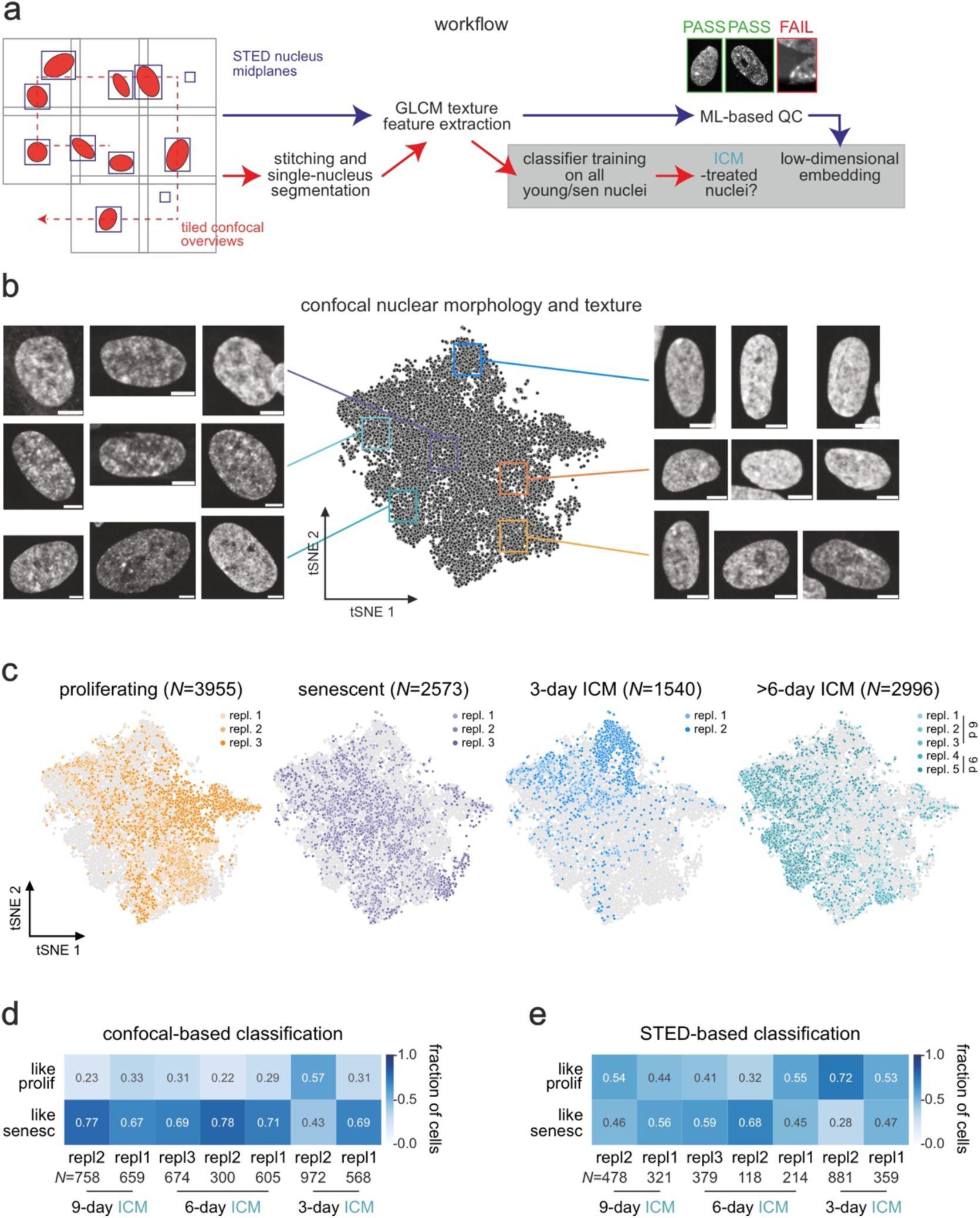
An automated classifier approach for assessing senescent cell nuclear morphology. **a.** Overview of the automated imaging and analysis workflow. Coarse-resolution confocal stacks were acquired in a tiled fashion and, once a nucleus of sufficient quality was detected, a mid-plane STED image of it was also acquired. GLCM features were extracted from single nuclei from confocal (after tile stitching and maximum projection of z-stacks) or STED images and used for downstream embedding and classification. **b.** t-SNE embedding of features extracted from confocal images with randomly-selected example images from proliferating (orange), senescent (purple), ad ICM-treated cells (turquoise) shown. Bar, 5 μm. **c.** As in panel B, but highlighting proliferating, senescent and 3-or 6-day ICM-treated cells of different replicates. **d.** Confusion matrix showing the fraction of ICM-treated cells classified as similar to proliferating or senescent cells using an SVM classifier trained on confocal nuclear features. **e.** An in panel D, but using a classifier trained on STED nuclear features.

We extracted GLCM texture (e.g., homogeneity, contrast, entropy and angular second moment) and other features (e.g., area, eccentricity, mean intensity) from each STED nuclear image that passed this QC, and from full nuclei in confocal images, and used them for t-SNE embedding (Fig 2b). In parallel, features from proliferating and replicatively senescent cells only were used to train a Support Vector Machine (SVM) classifier and assess the extent to which cells treated with ICM for different numbers of days resembled senescent ones. Using features extracted from confocal images, we recorded a broad range of nuclear phenotypes. t-SNE embedding showed that most proliferating cells separate from senescent ones, albeit with considerable mixing of replicates (Fig 2c); this is in line with previous single-cell transcriptional profiling that identified senescent-like cells in “young” populations and *vice versa*^20^. We also saw 3-day ICM-treated cells predominantly clustering away from senescent and proliferating ones (likely representing an intermediate state), whereas 6-and 9-day ICM-treated cells were markedly adjacent to senescent rather than 3-day or proliferating ones (Fig 2c). These degrees of separation also manifested in the SVM-based classification of ICM-treated cells, where 6-and 9-day-treated IMR90 classified as senescent, and 3-days cells scored as more ambiguous (Fig 2d). Notably, when STED images were used, sample-to-sample variation overshadowed phenotypic differences between proliferating and senescent nuclei resulting in the inconclusive classification of ICM-treated cells (Fig 2e). We attribute this to the large effects that even small variance in, for example, DNA staining intensity can have on fine scale details captured by STED nanoscopy. This was also observed in a deep learning-based study classifying senescence from images of nuclear-stained cells, whereby coarse features provided more predictive power than finer-scale ones^77^. Overall, we could deduce that ICM treatment of IMR90 produced nuclear features resembling those of senescent cells, but exhibiting far less heterogeneity (see replicate dispersal in Fig 2c).

### ICM-induced gene expression changes resemble replicative senescence

We followed up the phenotypic characterization with gene expression profiling of ICM-treated IMR90. First, based on previous data on RS, we would expect overall reduced RNA production levels if ICM-cells cells had indeed committed to senescence. We measured this by incorporating EUTP into nascent RNA with a short pulse and then visualizing labelled transcripts via an A488 fluorescent tag. Following quantification of signal intensity in the different cellular compartments, we saw an almost 2-fold drop in nuclear and nucleolar RNA levels by 6 days of ICM treatment, but only a modest decrease in labelled cytoplasmic RNA. This resembled the progressive drop seen in IMR90 passaged into senescence (Fig 3a).

**Fig 3.**
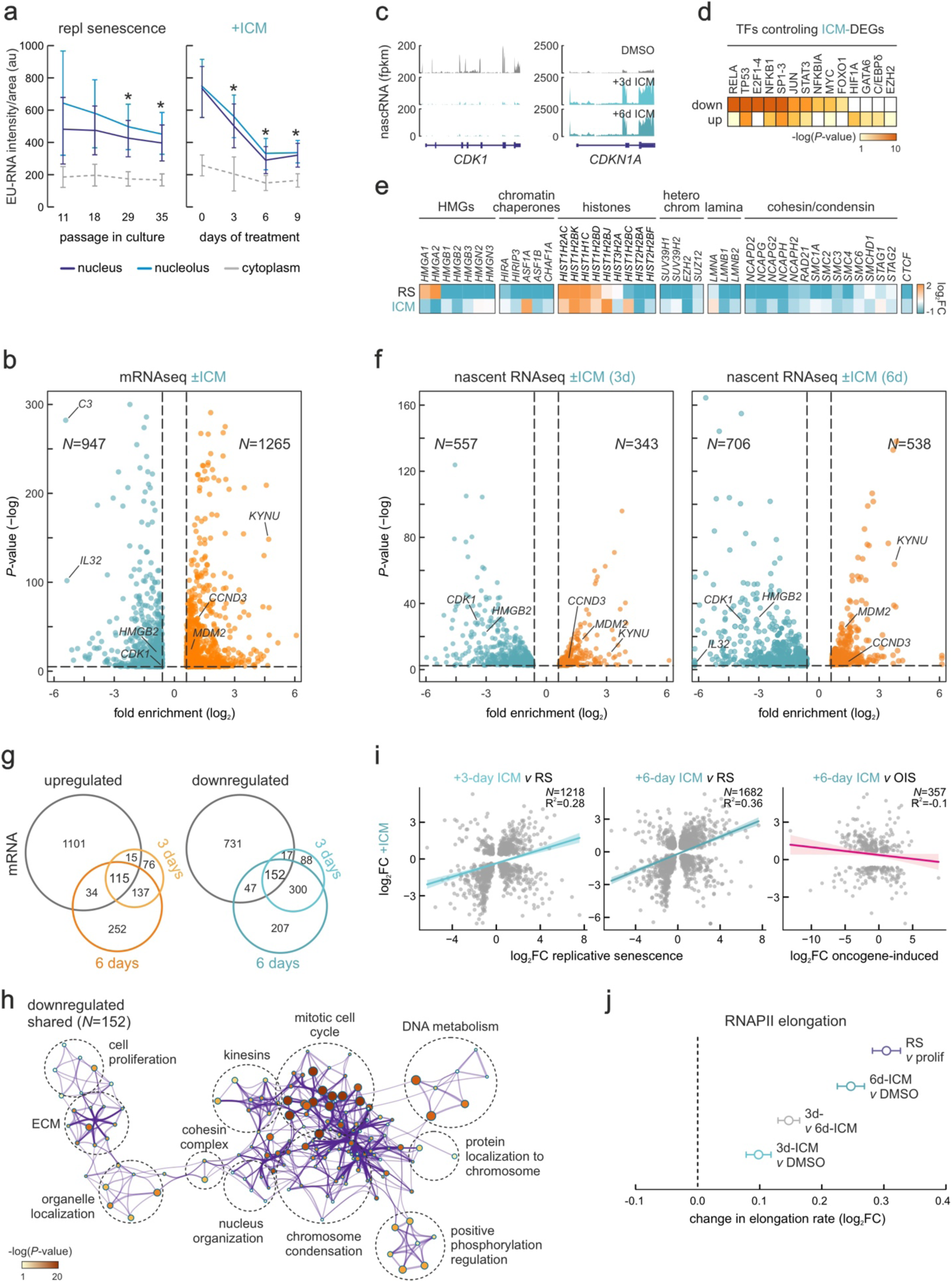
Transcriptional changes in ICM-treated human lung fibroblasts. **a.** Quantification of nascent EU-RNA levels (by immunofluorescence) in IMR90 passaged into senescence (left) or treated with ICM (right). *: significantly different to starting nuclear/nucleolar levels; *P*<0.05, unpaired twotailed Student’s t-test. **b.** Volcano plot showing all differentially expressed mRNAs between proliferating and ICM-treated IMR90. Significantly up-(orange; >0.6 log2FC) or down-regulated ones (green; <-0.6 log2FC) are indicated. **c.** RNA-seq profiles in the *CDK1* and *CDKN1A* locus from proliferating, 3-and 6-day ICM-treated IMR90s. **d.** Heatmap showing transcription factors predicted to regulate genes from panel C based on motif enrichment. **e.** Heatmap showing changes in mRNA levels upon senescence and ICM treatment of genes encoding selected chromatin-associated factors. For each gene shown, statistically significant expression changes (log2FC) were recorded in at least one condition. **f.** Volcano plot showing nascent RNA differences (fold enrichment) between 3 days and 6 days of ICM treatment. Significantly up-/downregulated genes are shown (>|0.6| log2-fold change). *N* is the number of the genes in each group. **g.** Venn diagrams of up-/down-regulated genes from ICM mRNA-seq and 3-or 6-day ICM nascent RNA-seq. **h.** GO term/pathway analysis of all commonly downregulated genes from panel G. **i.** Comparison of differentially expressed genes between replicative senescence and 3 or 6 days of ICM treatment (left and middle panel, R2=0.28 and 0.36, respectively) and oncogene induced senescence and ICM (right panel, R2=-0.1). *N* is the number of genes in each comparison. **j.** Plot showing changes (log2FC±SD) in mean RNAPII elongation rates calculated using nascent RNA-seq data.

Despite this documented drop, senescence entry is characterised by a distinct program involving both up-and downregulated genes^9,19,20^. We sequenced poly(A)^+^-selected RNA (mRNA-seq) from two different IMR90 isolates (I-10 and I-78) treated with DMSO (control) or ICM for 6 days. Following data analysis, we found ∼950 and >1250 genes to be significantly (*P*_adj_<0.05, log_2_FC>|0.6|) differentially expressed (Fig 3b,c). These genes could be linked to gene ontology (GO) terms associated with landmark senescence pathways, such as cell cycle regulation and growth factor response (for downregulated ones; Supplementary Fig 2a) or ECM reorganization and the p53 pathway (for upregulated ones; Supplementary Fig 2b). Surprisingly, and contrary to the prominent proinflammatory induction seen in senescence^19,44^, we found inflammatory activation via TNFα/NF-κB and interleukins to be markedly suppressed by ICM treatment, suggesting suppression of the SASP. This agreed with the anti-inflammatory mode-of-action of ICM in neurons^37^. Along these lines, prediction of transcription factors (TFs) that regulate these differentially expressed genes via TTRUST^45^ revealed an expected enrichment for p53 and E2F-family TFs for downregulated genes, but also strong enrichment of NF-κB subunits (RELA and NFKB1) and co-regulators (NFKBIA) involved in the inflammatory response (Fig 3d). Also, chromatin-associated gene markers known to be regulated upon RS entry were similarly affected by ICM treatment with the notable exception of HMGA1 and -A2, which have been implicated in the induction of heterochromatic foci in senescence and SASP regulation^14,46–48^ (Fig 3e).

Given that replicative senescence is a gene expression program predominantly regulated at the level of transcription^19^, we also sequenced and analysed nascent RNA profiles from 3-and 6-day ICM-treated IMR90 by using our “factory” RNA-seq approach^49^. Using the same cutoffs as before, but analyzing intronic RNA levels (reflecting direct transcriptional changes), we identified 343 and 557 up-and downregulated genes, respectively, at 3 days post-ICM treatment. These numbers increased to 538 and 706 after 6 days of treatment (Fig 3f). Despite this 25-55% increase, the majority of 3-day differentially expressed genes (i.e., 74% of up-and 81% of downregulated genes) were also identified at the 6-day mark (Fig 3g) in line with the high convergence between the two time points (Supplementary Fig 4c). Both sets associated with GO terms characteristic of RS induction (e.g., mitotic cell cycle, p53 pathway, telomere organization; Supplementary Figs 3a,b and 4a,b) and highly similar to those obtained following mRNA-seq analysis (Supplementary Fig 2). Similar were the TFs predicted to control the 3-and 6-day ICM-regulated genes, with p53 and E2F-family factors being most enriched (Supplementary Figs 3c and 4c). However, there was a relative de-enrichment for NF-κB-regulated downregulated genes at either time point, indicating that their suppression upon ICM treatment might not be exclusively transcriptional. Indeed, by looking at the overlap between differentially expressed mRNAs and nascent RNAs, only about 28% of up-or of downregulated nascent transcripts were also regulated at the messenger level, while >1,000 up-and >700 downregulated mRNAs did not qualify as differentially expressed in factory RNA-seq data (Fig 3g). Some part of this can be attributed to a difference in approach and analysis (exon-vs intron-level quantification), but likely also points to post-transcriptional regulation (e.g., changes in mRNA stability). Still the core of the 152 commonly downregulated genes in the three datasets concerned the expected types of GO terms and pathways (Fig 3h).

Next, we used publicly-available RNA-seq data from replicative ^50^ and oncogene-induced senescence in IMR90 ^9^, as well as a signature deduced from different types of senescence ^9^ for a comparison to data from 3-and 6-day ICM-treated cells. We found a robust positive correlation with RS differentially expressed genes (R^2^=0.28 and 0.36 for 3 and 6 days, respectively) and with the consensus senescence signature (R^2^=0.84; Supplementary Fig 5a), but no correlation with OIS ones (R^2^=-0.1; Fig 3i). This agrees with our phenotypic characterization showing the resemblance of ICM-induced senescence with replicative senescence, but a key discrepancy concerned inflammatory activation. Thus, we correlated RS and ICM mRNA-seq (where suppression of proinflammatory genes was most apparent; Supplementary Fig 2a) to validate this. The 175 genes strongly upregulated in RS, but suppressed by ICM treatment, were predominantly associated with the inflammatory response and the SASP (Supplementary Fig 5a,b).

Finally, we examined two measures of cellular ageing, the increase in RNAPII velocity in senescent cells^24^ and the changes in methylation levels at six senescence-predictive CpGs^51^. The former showed the expected acceleration of RNAPII (Fig 3j), while the latter showed no predictive power during a 12-day ICM treatment in contrast to how it performs for RS (Supplementary Fig 5c).

### Comparison of replicative and ICM-induced senescence at the single cell-level

One key motivation behind the pursuit of this system of ICM-triggered senescence was the need for a more synchronous and homogeneous induction of senescence in a given cell population, as the entry into RS is largely stochastic and heterogeneous at the level of individual cells (see ref.^20^ for RS in HUVECs). This is not only due to idiosyncratic cell-intrinsic properties (e.g., telomere attrition), but also a result of complex paracrine signaling via the SASP^44^. Thus, we reasoned that the apparently SASPless ICM phenotype would produce homogeneously-senescent cell populations within 6 days of treatment.

To test this, we generated single-cell transcriptomic data from proliferating (DMSO-treated), replicative senescent, and 6-day ICM-treated IMR90 from the same isolate. We interrogated a total of ∼26,000 cells (8,443 proliferating, 7,947 senescent, and 9,354 ICM-treated) using 3’ end single-cell RNA-seq. Following analysis of >1.8 billion reads (mean coverage was >70,000 reads/cell, median number of genes detected was 5,697 genes/cell), single-cell transcriptomes that met standard quality controls (Fig 4a,b) were used for in unsupervised clustering. This produced 5 clusters, as reflected in t-SNE embedding, of which one (cluster 1) was almost exclusively populated by proliferating and one (cluster 3) equally exclusively by ICM-treated cells (Fig 4c). In accordance to the notion of RS heterogeneity ^20^, some proliferating cells clustered among senescent cells (mostly in cluster 0) and some senescent cells mixed with proliferating ones (in cluster 1). However, ICM-treated cells mixed only with senescent ones (in clusters 0, 2, and 4) and showed overall less dispersion in t-SNE plots (Fig 4c). This was also reflected in the distribution of characteristic senescence markers, like *HMGB2*, *DNMT1*, and *CDKN1A*. *HMGB2* and *DNMT1* were expressed in proliferating cells of cluster 1 only and essentially not at all in senescent and ICM-treated cells alike, while *CDKN1A* expression was confined to senescent and ICM-treated cells, the latter showing significantly higher activation (Fig 4d,e). Largely uniform and low *CTCF* expression levels (as most transcription factors are more lowly expressed) provide a control (Fig 4d,e).

**Fig 4.**
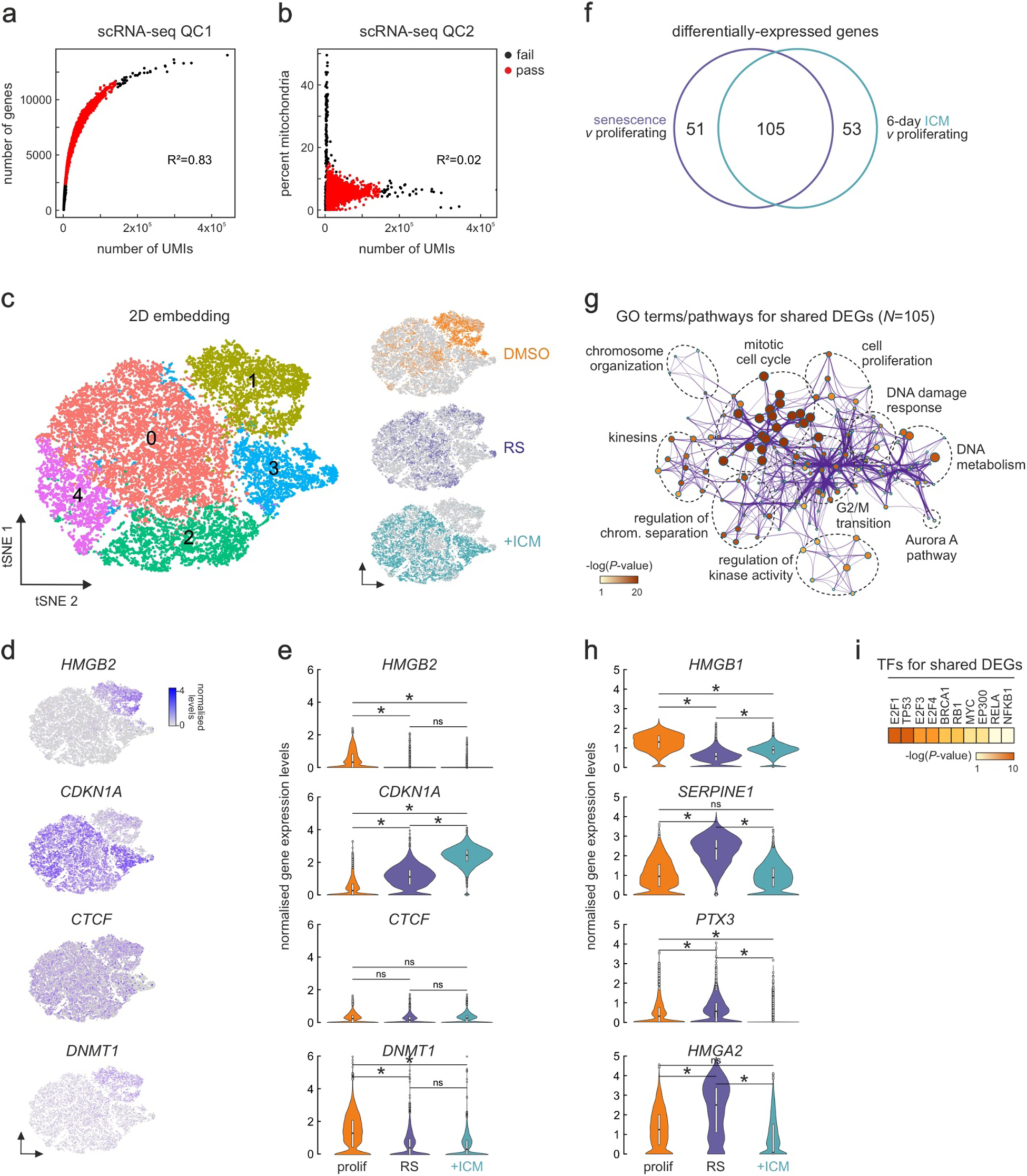
Single-cell analysis of ICM-induced transcriptomes. **a.** Scatter plot of the number of unique molecular identifiers (UMIs) versus the number of detected genes in each cell analyzed. Cells that passed (red) or not (black) this quality filter and the calculated Spearman’s correlation coefficient (R2) are indicated. **b.** As in panel a, but for the number of UMIs versus the percent of mitochondrial genes detected in each cell. **c.** Left: t-SNE embedding of gene expression profiles from 25,744 cells clustered in an unsupervised manner. Right: Projection of proliferating (DMSO), senescent (RS), and 6-day ICM-treated cells (+ICM) onto the t-SNE map. **d.** Projection of selected marker gene expression levels onto the t-SNE map of panel C. **e.** Violin plots showing expression level distribution of the marker genes from panel D in proliferating (prolif), senescent (RS), and 6-day ICM-treated cells (+ICM). *: *P*<0.01, Wilcoxon-Mann-Whitney test. **f.** Venn diagram showing the overlap of differentially expressed genes from senescent and ICM-treated cells. **g.** GO term/pathway analysis of the 105 shared differentially expressed genes from panel F. **h.** As in panel E, but for exemplary SASP-related genes. *: *P*<0.01, Wilcoxon-Mann-Whitney test. **i.** Heatmap showing transcription factors predicted to regulate genes from panel F based on motif enrichment.

Next, we identified condition-specific differentially expressed genes. Using a threshold of log_2_FC>|0.25| and comparing all proliferating with either senescent or ICM-treated cells, we detected >150 differentially expressed genes per condition. Of these, 2/3 (i.e., 105 genes; more than expected by chance, *P*<0.01) were shared between senescent and ICM-treated cells highlighting their convergence (Fig 4f) and associated with GO terms central to the senescent phenotype, like cell cycle regulation, chromosome organization, or DNA metabolism (Fig 4g). Notably, almost all of the 105 genes were included in the 267 commonly differentially regulated genes seen by bulk RNA-seq measurements (Fig 3h).

Last, we asked how SASP regulation manifested in this data. Looking among differentially expressed genes for markers derived from the SASP atlas (http://www.saspatlas.com), like *SERPINE1* and PTX3, or genes indirectly controlling SASP production like *HMGA2*^13^, we found these were significantly upregulated across senescent cells, but reduced to below control levels in ICM-treated IMR90 (Fig 4h). This was reflected in the de-enrichment for genes regulated by RELA and NFKB1 in the 105 differentially expressed genes shared by ICM-treated and senescent cells (Fig 4i) and agreed with observations from bulk RNA-seq data (Supplementary Fig 5a,b) and TTRUST analysis (Fig 3d). Thus, our single-cell analyses also confirmed the senescence-like features of ICM-induced cell growth arrest, as well as the more homogeneous nature of the response in IMR90 compared to senescence entry by continuous passaging.

### ICM-induced changes to the proteome are transcriptionally driven

Despite strong indications from our gene expression analyses about the convergence of replicative and ICM-induced senescence, it remained unclear whether the proteome also responded in the manner expected of senescent cells. To address this, we generated Ribo-seq and whole-proteome data from proliferating and ICM-treated IMR90 in biological triplicates, and compared them to equivalent data generated previously for RS entry^19^. Whole-proteome analysis after 6 days of ICM treatment revealed 565 significantly up-and 626 downregulated proteins (*P*<0.05, log_2_LFQ>|0.6|; Fig 5a). GO term analysis of these differentially expressed proteins identified senescence hallmark pathways linked to both up-(e.g., stress response and p53 pathway) and downregulated ones (e.g., cell cycle, chromosome organization, and RNA metabolism) (Fig 5b and Supplementary Fig 6a,b). This is in line with the differential analysis of RNA-seq data, as is the TTRUST query predicting that the genes coding for downregulated proteins were controlled by p53, MYC and E2F-family TFs (Supplementary Fig 6c).

**Fig 5.**
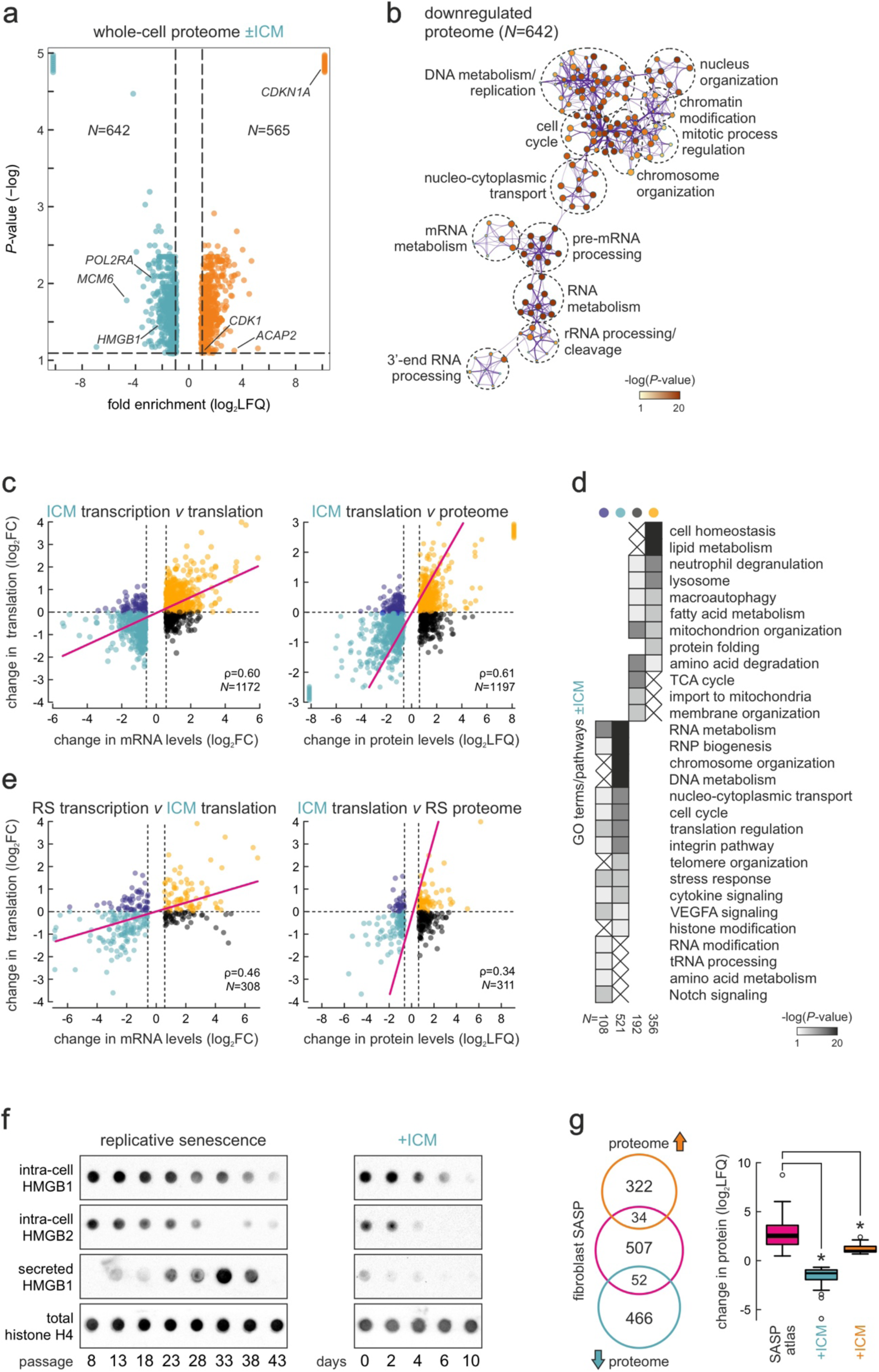
Proteomic changes induced by ICM treatment of IMR90. **a.** Volcano plot showing whole proteome up-(orange, >0.6 log2LFQ) and downregulated proteins (turquoise, <-0.6 log2LFQ) upon 6 days of ICM treatment. The number of proteins (*N*) in each set is indicated. **b.** GO term/pathways analysis of all downregulated proteins from panel A. **c.** Left: Scatter plots showing correlation between mRNA-seq (transcription) and Ribo-seq levels (translation) of transcripts differentially expressed upon 6-day ICM treatment. Right: Correlation between mRNA-seq and whole proteome levels for the same set of genes. The number of genes/proteins (*N*) in each set and Pearson’s correlation coefficient (ρ) are indicated. **d.** Heatmap showing GO terms/pathways associated with transcripts in the different quadrants of panel C (colorcoded the same way). The number of transcripts in each subgroup (*N*) is indicated. **e.** As in panel C, but using differentially expressed genes from replicative senescence. **f.** Dot blot showing intracellular HMGB1 and HMGB2 and secreted HMGB1 levels across passages (left panel) and days of ICM treatment. Histone H4 levels provide a control. **g.** Left: Venn diagram showing up-and downregulated proteins from whole-cell proteomics crossed with all known fibroblast SASP factors. Right: Box plot showing changes in SASP-related protein levels (log2LFQ). *: *P*<0.01, unpaired two-tailed Student’s t-test.

We next focused on the regulation of translation upon ICM treatment. We previously used Ribo-seq to show that almost none of the changes in senescence-related gene expression could be explained by changes in translation levels only^19^, and that no ribosome stalling, competition by upstream ORFs or translational deficiency were found^52^. Thus, we generated Ribo-seq data for 6-day ICM-treated IMR90 and correlated them to matching mRNA-seq and whole-cell proteome datasets. Much like what we observed for RS, all significant changes in translation were explained by an equal change in transcript availability (Fig 5c). These transcripts were linked to key processes for senescence commitment like the downregulation of RNA metabolism, cell cycle regulation, and telomere organization (Fig 5d). There were 300 transcripts “buffered” by translation (i.e. transcriptionally suppressed but translationally boosted or *vice versa*), which could be implicated in pathways like RNA modification or Notch and VEGFA signaling (Supplementary Fig 5c,d). These correlations remained largely unchanged when mRNA-seq or Ribo-seq data were replaced by those generated in RS IMR90 (Fig 5e). Thus, the ICM-induced gene expression program is predominantly regulated at the level of transcription, just like the one of RS^19^.

Interestingly, and in line with our mRNA-seq analysis, the expression of proteins involved in cytokine stimulation and the interferon response was suppressed by ICM treatment (Fig 5A and Supplementary Fig 6a), and these transcripts were mostly found “buffered” when Ribo-seq was co-considered (Fig 5d). Dot bot analysis of intracellular and extracellular levels of HMGB1 and -B2 showed that, in contrast to what was observed during extended IMR90 passaging, HMGB1 is not released into the growth media to act as a proinflammatory “alarmin”^53^ despite its apparent intracellular reduction (Fig 5f). We also assessed this more broadly by using a public catalogue of fibroblast SASP (http://www.saspatlas.com) and overlapping it with ICM-induced proteome changes. Of ∼600 *bona fide* SASP factors, <15% we overlapped our data, with 52 being significantly down-and only 34 significantly upregulated (Fig 5g). However, even those upregulated by ICM, were not induced to the full extent observed in senescent fibroblasts (Fig 5g). Taken together, the above analyses confirm that ICM triggers a SASPless replicative-like senescent phenotype.

### ICM triggers 3D genome reorganization reminiscent of RS

RS has been linked to extensive reorganization of 3D chromatin folding^54^, with entry into senescence already characterised by changes at the level of compartments and TADs^14,20^. Therefore, as a last element in the characterization of ICM-induced senescence, we addressed the extent of 3D genome reorganization after 6 days of treatment. We generated high-resolution Micro-C data^55^ from proliferating (DMSO-treated) and ICM-treated IMR90. Replicates form each condition were sequenced to >1.1 billion read pairs generating maps with >614 and >711 million contact pairs for proliferating and ICM-treated cells, respectively. Of these, >50% represented long-range (separated by >10 kbp) interactions (see Supplementary Table 1 for details). As a result, we obtained dense 5-kbp resolution contact maps with differences between conditions (Fig 6a).

In more detail, interaction decay plots showed reduced contact frequency at the Mbp scale, in parallel with increased frequency at sub-Mbp separation distances (Fig 6b). The former is reminiscent of a “better compartment definition” we previously observed in lower resolution Hi-C data from senescent IMR90 and HUVECs^20^, and was corroborated by reduced inter-compartmental interactions in ICM-treated cells (Fig 6c). The latter can be explained as an effect associated with ICM-induced suppression of transcriptional output (Fig 3a) leading to chromatin compaction at the sub-TAD level^56^.

**Fig 6.**
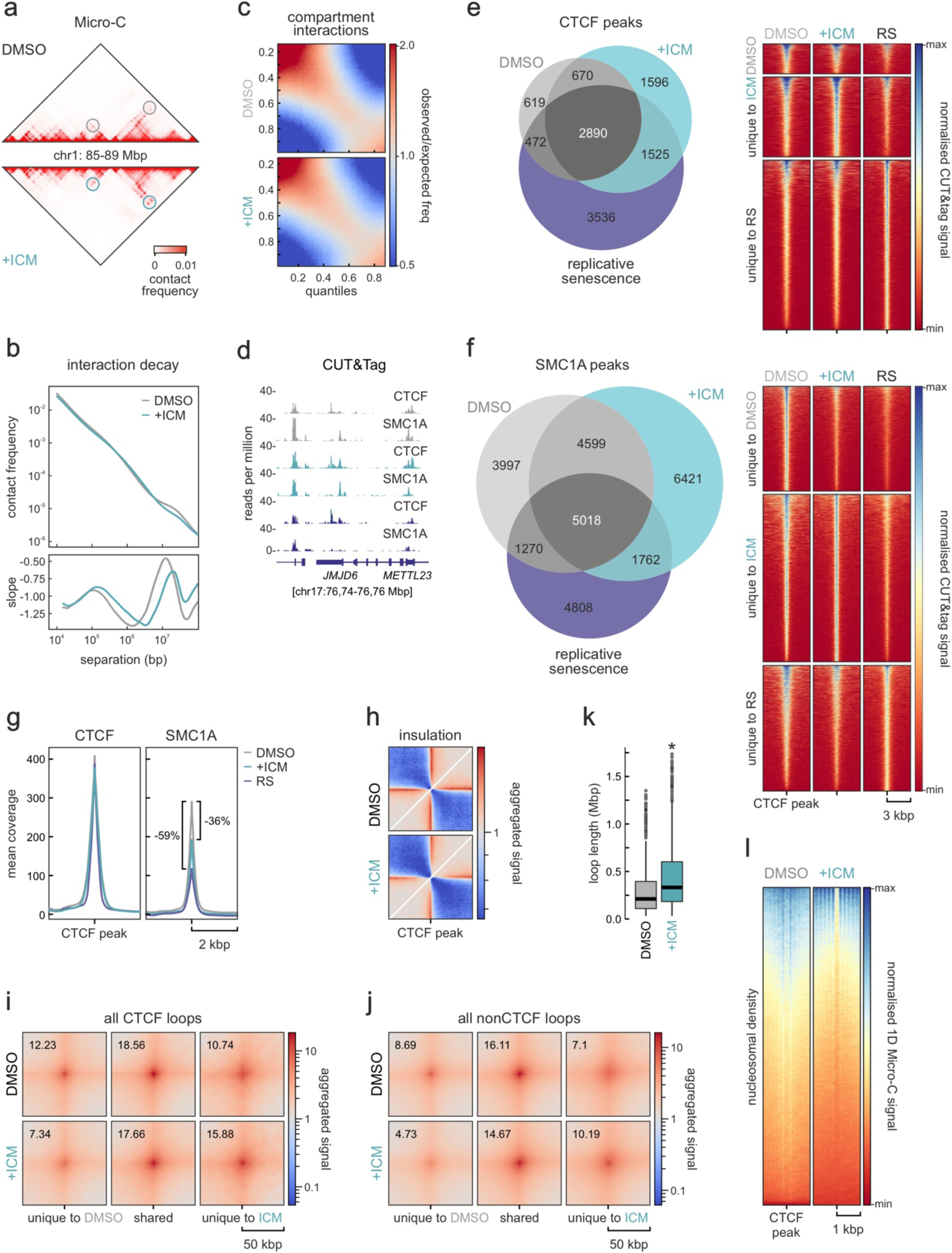
3D genome reorganization at 6 days post-ICM treatment. **a.** Heatmap showing Micro-C contacts at 10-kbp in an exemplary 4-Mbp segments from chr1. Differences in loop formation between proliferating (DMSO) and ICM-treated IMR90 (+ICM) are denoted (circles). **b.** Plots showing decay of contact frequency as a function of genomic distance (top) and its first derivative (bottom) for proliferating and ICM-treated cells. **c.** Saddle plots showing contact distribution among and between inactive (top left corner) and active compartments (bottom right corner) in proliferating (DMSO) and ICM-treated Micro-C data (+ICM). **d.** Representative genome browser views of CTCF and SMC1A CUT&Tag signal along a 100-kbpp region of chr17 from proliferating (grey), ICM-treated (green) or senescent IMR90 (purple). **e.** Left: Venn diagram showing the overlap of CTCF peaks (top 1%) in CUT&tag data from proliferating, ICM-treated, and senescent IMR90. Right: Heatmaps showing scaled CUT&tag signal in the 4 kbp around the peaks. **f.** As in panel E, but for SMC1A CUT&Tag data. **g.** Line plots showing mean CTCF or SMC1A CUT&Tag signal coverage in the 4 kbp around all CTCF peaks from proliferating (grey), ICM-treated (green) or senescent IMR90 (purple). **h.** Insulation plot averaging Micro-C contacts in the 600 kbp around CTCF peaks from proliferating (DMSO) and ICM-treated IMR90 (+ICM). Aggregate plots showing average Micro-C signal in the 100 kbp around CTCF loop summits called at at 5-kbp resolution from unique to or shared by proliferating and ICM-treated IMR90. **j.** As in panel I, but for nonCTCF-anchored loops. **k.** Box plots showing the distribution of CTCF loop lengths in proliferating (DMSO) and ICM-treated IMR90 (+ICM). *: *P*<0.01, Wilcoxon-Mann-Whitney test. **l.** Heatmaps showing nucleosome distribution signal derived from Micro-C data in the 2 kbp around CTCF motifs under CUT&Tag peaks from proliferating (DMSO) or ICM-treated IMR90 (+ICM).

We also generated CUT&Tag data for the two key architectural factors giving rise to chromatin loops (Fig 6a), the insulator protein CTCF and the ring-shaped complex of cohesin via its SMC1A subunit^57^. Analysis of CTCF CUT&Tag returned 4,652 CTCF peaks in proliferating, 8,440 in senescent, and 6,666 in ICM-treated IMR90 (using the top 1% of peaks and filtering for the presence of a consensus CTCF motif under each peak; see Methods) (Fig 6d,e). Similarly, 14,884 SMC1A peaks were called in proliferating, 12,928 in senescent, and 17,810 in ICM-treated cells (Fig 6d,f). Overall, ICM CTCF peaks overlapped more peaks from senescent rather than proliferating IMR90, but the converse was true for SMC1A peaks (Fig 6e,f). Nevertheless, inspection of signal distribution in genome browser tracks and in heatmaps showed that there were indeed numerous strong CTCF peaks emerging in both ICM-induced and replicative senescence, while for SMC1A this mostly held true in ICM-treated cells (Fig 6d-f). However, SMC1A signal at shared CTCF peaks was reduced by at least 36% in ICM-induced as well as in RS cells (Fig 6g).

The emergence of condition-specific CTCF-and cohesin-occupied positions along IMR90 chromosomes in combination with reduced cohesin levels led to a decrease in overall contact insulation around CTCF peaks (Fig 6h). This effect should translate into formation of condition-specific loops from the 22,871 and 14,995 called in proliferating and ICM-treated IMR90, respectively. After stratifying loops as CTCF-or nonCTCF-anchored, we indeed found 613 CTCF (out of 1,801; 34%) and 4,303 nonCTCF loops (out of 13,194; 33%) that were unique to ICM-treated cells and showed increased contact frequency (Fig 6i,j). Notably, and despite the fewer loops called in ICM data, loop length was significantly increased following ICM treatment (Fig 6k). This agreed with previous observations from Hi-C and CTCF HiChIP data in RS^20^, and may be linked to the formation of senescence-induced CTCF clusters (SICCs). Last, we exploited the fact that Micro-C signal contains information about nucleosome positioning^55^ to examine their density genome-wide. We plotted 1-D Micro-C signal around CTCF peaks and found that, despite an overall signal decrease in ICM-induced cells, nucleosomes were positioned in better defined arrays following ICM treatment (Fig 6l). Nucleosome signal decrease is characteristic of senescence^24^, while the more defined positioning is reminiscent of what was seen following depletion of RNAPII in human cells^58^. Thus, also as regards 3D genome reorganization, ICM-induced senescence aligns well with RS entry.

## Discussion

Replicative senescence is essentially hard-wired in the homeostatic program of proliferating cells grown *in vivo* and *in vitro*^38^. However, its emergence in cell populations is non-homogeneous and heavily influenced by various waves of paracrine signaling^34,59^. This poses several issues when studying cell commitment to senescence. For example, the core transcriptional program responsible for senescence entry might be obscured by differential gene induction and silencing due to SASP and secondary Notch signaling in a cell population. Similarly, studying the temporal order of molecular events leading up to senescence commitment can be obscured by the non-synchronous manner by which it occurs in different cells. Finally, the recent interest in senolytic^60,61^, senogenic^62,63^, and partial reprogramming regimes applied to aged cells ^64,65^ could also benefit from a controlled *in vitro* model for screening and/or deep functional characterization of candidate compounds.

Here, we present a phenotypic and multi-omics characterization of senescence induced in the prototypic IMR90 cellular model following treatment with the small molecule inhibitor ICM. We showed that, within 6 days of treatment with 10 μM ICM, replicative-like senescence is stably and irreversibly instated in the cell population in a synchronous and homogeneous manner. Most notably, senescence emerges in the absence of apparent paracrine signaling by the SASP that turns on proinflammatory genes^44^. However, at first glance this seems to contrast the original characterization of the ICM mode-of-action^37^. On the one hand, ICM was indeed developed and selected for its robust anti-inflammatory capacity. This stemmed from its presumed ability to constrain the release of HMGB proteins, especially HMGB1, from cell nuclei by interfering with their post-translational modification^37^. HMGB1 has a pronounced role as an “alarmin” released from cells to trigger inflammation in its niche *in vitro* and *in vivo*^18,53,66,67^. ICM treatment constrains neuroinflammation in cell cultures and animal models^37^ and HMGB1-related secretion and signaling from pancreatic cells^68^ much like it constrains the SASP in IMR90 in our hands. However, the compound was never tested using treatment periods longer than 24 h, which would have allowed its full effects to deploy. This would then be the synchronous and near-complete nuclear depletion of both HMGB1 and HMGB2 from cell nuclei, but without their apparent rerouting to the secretory pathway (although HMGB2 is not secreted by senescent IMR90, but rather turned over^19^). Thereby, ICM cannot permanently restrict HMGBs in the nucleus, but can block secretion in favour of degradation, likely via the crosstalk of HMGB1 with the autophagic pathway^69^. It would, thus, be interesting to eventually examine multi-tissue effects following systematic *in vivo* ICM administration in animal models in more detail, while factoring in our observations from IMR90.

In summary, we hoped to share ICM-induced senescence with the broader community as a new tool for accelerating research into the core senescent program and the different components that mediate its irreversible nature (or allow for its sporadic bypass). Such rapid and synchronous pharmacological induction of senescence, devoid of any paracrine influence, could find various applications in the study of anti-ageing and late-life disease solutions.

## Acknowledgements

We would like to thank members of all laboratories involved in this study for helpful discussions. We thank the MPI-NAT Facility for Synthetic Chemistry for synthesizing the 610CP-C6-ICM compound, the Center for Advanced Light Microscopy (CALM) at LMU Munich for automated imaging of nuclear phenotypes, and the Cologne Center for Genomics for library preparation and high throughput sequencing. This work was supported by the German Research Foundation (DFG) via TRR81 (Project 109546710 awarded to AP), the SPP2202 Priority program (Projects 422389065 to AP and 422857584 to HH and HL), the KFO5002 Clinical Research Unit (Project 426671079 to AP), as well as by the SFB1565 (Project 469281184 to AP). SP, KS, NJ, ADM, and VVM were also supported by the International Max Planck Research School for Genome Science, YZ by the International Max Planck Research School for Molecular Biology, and AS by Erasmus+ funds.

## Author contributions

SP, KS, NJ, and AZ performed experiments; AS, VVM, YZ, NJ, and CN performed computational analyses; ADM analysed single-cell genomics data; AOC, GL, JK, and AM generated and analysed Ribo-seq data; DH, SN, HH, and HL generated imaging data and devised the classification pipeline; GL produced the 610CP-C6-ICM compound; AP and AB provided the RNAPII speed calculations; WW provided methylation data; AP conceived the study; SP and AP wrote the manuscript with input from all co-authors.

## Methods

### Cell culture and senescent assays

Single IMR90 isolates (I90-83, passage 5; Coriell Biorepository) were continuously passaged at 37°C under 5% CO2 in Minimal Essential Medium L-Glutamine without HEPES (MEM 1X) (Gibco™ Life Technologies GmbH, 31095052) supplemented with 10 % FBS (Life Technologies, 10500064), 1X (1%) MEM Non-essential Amino Acid Solution without L-glutamine (Sigma-Aldrich, M7145-100ML) and 1% Penicillin/Streptomycin (Gibco™ Life Technologies, 15140122). The senescent state of the cells was addressed by senescence-associated β-galactosidase assay (Cell Signaling) according to the manufacturer’s instructions. Cells were driven into senescence either by continuously passaging them to replicative exhaustion or by using Inflachromene (concentration and period of treatment is depicted on individuals experiments). Cell proliferation was monitored using the Sartorius IncuCyte S3 Live-Cell Analysis System and acquiring a picture every 8h for a total of 11 days. Finally, DNA methylation at six selected CpG islands was measured by isolating genomic DNA at the different cell states and performing targeted pyrosequencing (Cygenia GmbH) as previously detailed^51^.

### Immunofluorescence and image analysis

Cells treated with ICM-610CP were cultured on coverslips for 3 days and DNA was subsequently stained with 5-SiR-Hoechst^70^ and fixed via incubation with 4% paraformaldehyde (PFA) in Dulbecco’s Phosphate-Buffered Saline (DPBS). For every other staining, cells grown on coverslips were fixed via incubation with 4% PFA in DPBS at RT for 10min and then permeabilized with 0,5% Triton-X in PBS for 10min. Blocking was performed with 1% Bovine Serum Albumin (BSA) in PBS at RT for 1h. Cells were then incubated with the primary antibody (diluted in 0.5% BSA/PBS) at RT for 1h at the indicated dilution: mouse monoclonal anti-HMGB1 (1:1,000; Abcam ab190377-1F3); rabbit polyclonal anti-HMGB2 (1:1,000; Abcam ab67282); rabbit polyclonal anti-CTCF (1:500; Active motif 61311); rabbit polyclonal anti-H3K27me3 (1:1,000; Diagenode C15410069); rabbit polyclonal anti-p21 (1:500; Abcam EPR362 - ab109520). The primary antibody was washed with PBS twice for 5min per wash. Cells were incubated with the secondary antibody (diluted in 0.5% BSA/PBS) at RT, in the dark for 1h at the indicated dilution: anti-rabbit Alexa488 (1:1,000, Abcam ab150077); anti-mouse Cy3 (1:1,000, Abcam ab97035). Cells were then washed with PBS twice for 5min per wash. ProLongTM Gold antifade reagent with DAPI (#P36931) was added to the cells. For visualizing nascent transcripts, cells were pre-incubated with 2.5mM 5-ethynyl uridine (EU) for 40min at 37°C in their growth medium, fixed and processed with the Click-iT EdU chemistry kit (Thermo Fisher). For image acquisition, a widefield Leica DMI8 with an HCX PL APO 63x/1.40 (Oil) objective was used. The acquired images were subsequently analysed with the FIJI software^71^. Measurements of nuclear immunofluorescence signal were generated using a mask drawn on DAPI staining to define nuclear bounds. Background subtractions were then implemented to precisely determine the mean intensity per area of each immune-detected protein.

### Automated cell imaging and feature classification

50-70K IMR90 cells were seeded onto coverslips in 6-well plates and left to grow at 37 ◦C, with 5% CO_2_ in MEM (M4655), supplemented with 1% Penicillin/Streptomycin (Pen/Strep, P4333) and 1% non-essential amino acids (M7145) all from Sigma Aldrich, and 10% fetal bovine serum (FBS, F7524, LOT: BCBX5319) for 24h before fixation and staining. For ICM-treated cells, we exchanged the medium to one supplemented with 7.5µM ICM after 24h and left them to grow for 3, 6 or 9 days before fixation, exchanging the medium daily. The cells treated with ICM for 6 and 9 days were split once and twice, respectively. Cells were then prepared for immunofluorescence as described above, but also incubated in a SiR-DNA^72^ staining solution (2 µM in PBS-T) for 90 min at room temperature. Afterwards, cells were rinsed and washed twice for 5 min with PBST. Finally, coverslips with cells were mounted onto glass slides in MOWIOL. 30 min after mounting, coverslips were sealed with clear nail varnish and dried for 20 min at room temperature.

We acquired confocal and STED images on an on a 3D STED microscope system from Abberior Instruments (Göttingen, Germany) using a 100x UPlanSApo 1.4 NA oil immersion objective (Olympus, Japan) and pulsed 640nm excitation and 775 nm depletion lasers. To acquire a large number of super resolution images in an unbiased fashion, we automated the operation of the microscope using the *specpy* Python interface to the Imspector microscope control software. In our automation pipeline, we first continuously imaged confocal overview stacks with 20% overlap in a spiral. Following the acquisition of each overview, we detected nuclei in the maximum intensity projection of the tile and adjacent tiles (stitched based on their stage coordinates) via unsupervised clustering of pixels based on their intensities and Gaussian and Sobel filter responses using k-Means followed by binary erosion of radius=3 to remove small background detections. For all connected foreground regions, we acquired a STED image of the middle z-plane before continuing with the next overview tile. All acquired images as well as microscope metadata were saved in custom HDF5-based files during the automated acquisitions. Using this pipeline, we could run the microscope unsupervised for prolonged periods of time and we typically imaged each sample for 12-24h.

To extract features from STED images, we proceeded as follows: For each STED detail image, we first normalized the intensities to the 0.025 and 0.995 quantiles and then performed a simple segmentation by Li thresholding on a strongly blurred (Gaussian blur with **σ** = 16**σ**) version of the image followed by removing small objects <512 px^2^ and filling holes smaller than 512 px^2^. Within the foreground area we calculated grey level co-occurrence matrices (GLCMs) at distances **∈ {2, 4, 7, 12, 16} and angles ∈ {0, π/2}** and calculated all summary statistics available in scikit-image’s greycoprops function (’contrast’, ‘dissimilarity’, ‘homogeneity’, ‘energy’, ‘correlation’ and ‘ASM’) from slightly blurred (**σ** = 0.5**σ**) versions of the normalized images. In addition, we calculated the mean foreground intensity in both the original and normalized image, the standard deviation of the foreground intensity, the number of pixels of the segmented area, the low and high quantiles of raw image intensities used in normalization, as well as the image width and height and number of rows and columns composed wholly of zeroes (an indication of images acquired outside the scanner’s field of view or shutdown of the detector due to too high light exposure, which we aimed to remove in a subsequent quality control step). To distinguish good STED images from erroneous detections during the automated imaging we used machine learning-based quality control. We manually sorted 493 images as good (i.e., containing a single complete and in-focus nucleus) or bad and trained a Random Forest classifier on their features (normalized to zero mean and unit variance). Using 5-fold cross-validation, we determined the probability threshold for the good class necessary to achieve 95% precision (true positive rate) and applied the classifier with this threshold to all uncategorized images. For the subsequent steps, we only used the images classified as good.

We then used the features of each cell (except auxiliary ones like the number of blank rows/columns or image size) and performed a 2-dimensional embedding using t-SNE. Furthermore, we used the features of all young and senescent samples to train a SVM classifier and applied it to all treated samples to see whether they would be preferentially classified as young or senescent. However, in both approaches, we arrived at inconclusive results and saw a strong dependence on individual replicates. We reasoned that at the fine scale captured by STED nanoscopy, small changes, e.g., in SiR-DNA staining intensity, have a stronger effect on nuclear texture than senescence state. The code for our image analysis pipeline was implemented in Python using the numpy/scipy^73^, scikit-image^74^ and scikit-learn^75^ libraries and is freely available at: https://github.com/hoerlteam/chromatin-texture-senescence/. Due to sample-to-sample variation seems to overshadow biological effects on the nanoscale in STED images, we decided to also analyze the confocal overview images we initially acquired as auxiliary data during the automated imaging. We extracted individual overview tiles from the combined HDF5 files, saved them as TIFF stacks and used BigStitcher^76^ to stitch (refining the stage positions recorded by the microscope) and fused them into one volume per acquisition run. We then used Cellpose to detect individual nuclei in a z-maximum projection and calculated GLCM texture features and summary statistics (with distances **∈ {2, 4, 8, 16*px*** and angles **∈ {0, π/2})** as well as shape and intensity features (area, eccentricity and mean intensity) in the z-projection for each detected nucleus individually. We normalized intensities for each image to the (0.025, 0.998) quantiles and applied a small amount of Gaussian blur (**σ** = 0.5**σ**) before extracting features. We then proceeded to perform t-SNE embedding as well as SVM-based classification of ICM-treated cells based on young and old samples as described for the STED data above.

### RNA isolation, sequencing and analysis

Proliferating, senescent and ICM-treated IMR90s were harvested in TRIzol LS (Life Technologies) and total RNA was isolated and DNase I-treated using the DirectZol RNA miniprep kit (Zymo Research). Following selection on poly(dT) beads, barcoded cDNA libraries were generated using the TruSeq RNA library Kit (Illumina) and were paired-end sequenced to >50 million read pairs on a HiSeq4000 platform (Illumina). Default settings of STAR aligner^78^ were used to map the raw reads to human reference genome (hg19) and quantification of unique counts was done via *featureCounts*^79^. The RUVs function of RUVseq^80^ was used to further normalize the counts, prior to differential gene expression estimation using DESeq2^81^. Genes with an FDR < 0.01 and an absolute (log_2_) fold change of > 0.6 were deemed as differentially expressed. GO term enrichment plots were generated using Metascape (http://metascape.org/gp/index.html)^82^. For RNA that was later used for qPCR the isolation procedure was the same as the one described above. cDNA was synthesized with SuperScript™ II Reverse Transcriptase (Invitrogen™ Life Technologies, 18064071) and random primers (Sigma-Aldrich, 11034731001) according to the manufacturer’s protocol. Full list of primers used for qPCR can be found in Supplementary Table 2. Finally, for analysis of nascent RNA in IMR90 the ‘‘factory RNA-seq‘‘ approach was applied on 5mil ICM-treated cells^83^, RNA was isolated and sequenced as above, and intronic read counts were obtained and differentially analyzed for the two conditions using the iRNAseq package^84^.

For RNAPII elongation rates calculated from “factory” RNA-seq data, annotation files were downloaded from Ensembl (https://www.ensembl.org/; version hg19). The following filtering steps were applied on the intronic ENSEMBL annotation files. First, we removed overlapping regions between introns and exons to avoid confounding signals due to variation in splicing or transcription initiation and termination. Overlapping introns were merged to remove duplicated regions from the analysis. In the next step, we used STAR to detect splice junctions and compared them with the intronic regions. Introns with at least five split reads bridging the intron (that is, mapping to the flanking exons) per condition were kept for subsequent analyses. When splice junctions were detected within introns, we further subdivided those introns accordingly. The slope of the intronic coverage was calculated in these introns across all samples as described^24^. To avoid artefacts due to the different numbers of introns used per sample, we always contrasted the same sets of introns for each comparison of different conditions, only using slopes that were negative across all samples. The Wilcoxon signed-rank test with continuity correction was used for statistical testing in R.

### Chromatin immunoprecipitation and qPCR

Proliferating and ICM-treated IMRO90s were cultured to 80% confluence in 15-cm plates and they were crosslinked in 15 mM EGS/PBS (ethylene glycol bis(succinimidyl succinate); Thermo) for 20 min at room temperature, followed by fixation for 40min at 4°C in 1% PFA. Cells were then processed with the ChIP-IT High Sensitivity kit (Active motif) according to the manufacturer’s instructions. Chromatin was sheared to 200-500 bp fragments via sonication using a Bioruptor Plus (25 cycles, 30 sec *on*/ 30 sec *off*, high input), immunoprecipitation was done using 4 μg of anti-HMGB2 antibody (Abcam ab67282) to approx. 30 μg of chromatin and the samples were incubated overnight in a rotor at 4°C. DNA was captured on protein A/G agarose beads and purified using the ChIP DNA Clean & Concentrator kit (Zymo) and used for qPCR. Oligos used in qPCR are listed in Supplementary Table 3.

### Ribo-seq and data analysis

High-throughput ribosome profiling (Ribo-seq) on proliferating, senescent and ICM-treated IMR90s was performed in collaboration with EIRNA Bio Ltd (https://eirnabio.com) according to their established protocol^85^. Three independent replicas of proliferating, senescent or ICM-treated IMR90 were grown, harvested in ice-cold polysome isolation buffer supplemented with cycloheximide, and shipped to Ribomaps for further processing and library preparation. Roughly 15% of each lysate was kept for RNA isolation and used for RNA-seq of poly(A)-enriched fractions on a HiSeq2500 platform (Illumina). After sequencing of both Ribo-and mRNA-seq libraries, the per base sequencing quality of each replicate passed the quality threshold, raw read counts were assigned to each protein-coding open reading frame (CDS) for Ribo-seq and to each transcript for mRNA-seq, and replicate correlations were tested. Read length distribution for Ribo-seq datasets fell within the expected range (25-35nt), showing strong periodic signals and an enrichment in annotated CDSs. For mRNA-seq, read lengths ranged between 47 and 51 nt and distributed uniformly across transcripts. For differential gene expression analysis, anota2seq^86^ was used. Changes in Ribo-seq data depict changes in the ribosome occupancy of the annotated protein-coding CDS, and thus, only ribosome-protected fragments that map to the CDS were used in the analysis. VST normalized counts outputted using DESeq2^81^ and inputted into anota2seq were used for all subsequent downstream analysis. Differences in genes that pass a default FDR threshold of 15% were considered regulated. Such significant differences are then categorized into one of the following three modes: (i) translational: Changes in Ribo-seq that are not explained by changes in RNA-seq and imply changes at the protein level are due to changes at the translational level; (ii) mRNA abundance: Matching changes in RNA-seq and Ribo-seq that infer changes at the protein level are predominantly induced by changes at the transcriptional level; (iii) *buffering*: changes in RNA-seq that are not explained by changes in Ribo-seq and suggest maintenance of constant protein levels induced by changes at the transcriptional level or *vice versa*.

### Protein extraction, western blotting, and mass spectrometry

Proliferating and ICM-treated IMR90s (approx. 2×10^6^ per condition) were gently scraped off 15-cm dishes. Cells were then pelleted for 5min at 1,200rpm. The supernatant was discarded and pellets were lysed in 150u of RIPA lysis buffer (20 mM Tris-HCl pH 7.5, 150 mM NaCl, 1 mM EDTA pH 8.0, 1 mM EGTA pH 8.0, 1% NP-40, 1% sodium deoxycholate) containing 1x protease inhibitor cocktail (Roche) for 30min on ice. Sample were then sonicated in low input for 3 cycles (30sec on/30sec off) and centrifuged for 15min at >15,000g. Then the supernatant was collected and the protein concentration was measured using the Pierce BCA Protein Assay Kit (Thermo Fisher Scientific). Rabbit polyclonal anti-HMGB2 (1:1,000; Abcam ab67282); rabbit polyclonal anti-CTCF (1:500; Active motif 61311); rabbit polyclonal anti-EZH2 (1:500; Active motif 39901); rabbit polyclonal anti-H3 (1:500; Abcam ab1791); mouse monoclonal anti-a tubulin (1:1000, Abcam ab7291) were used for blotting. For whole-cell proteomics, protein extracts in RIPA buffer were analyzed by the CECAD proteomic core facility in biological triplicates on a Q-Exactive Plus Orbitrap platform (Thermo Scientific) coupled to an EASY nLc 1000 UPLC system with column lengths of up to 50 cm. All proteins discovered via whole-cell mass spectrometry are listed in Supplementary Table 4.

### Cleavage Under Targets and tagmentation

0.5 million cells were lifted from plates using accutase, fixed with 0.3% PFA/PBS for 2min at RT and then quenched with 0.125M ice cold glycine for 5min at RT. Samples were then processed according to manufacturer’s instructions (Active Motif). Samples were paired-end sequenced to obtain more than 5×10^6^ reads, which were then processed exactly as described in https://yezhengstat.github.io/CUTTag_tutorial/. Briefly, paired-end reads were trimmed for adapter removal and mapped to human (hg38) and E. coli reference genomes (ASM584v2) using Bowtie 2 ^87^. E. coli mapped reads were then quantified and used for calibrating human-mapped reads. Peak calling was performed using a multi-FDR-tryout method (FDR <0.01 to <0.1). For CTCF and SMC1A, an FDR <0.01 was selected and only CTCF peaks with a canonical CTCF motif were considered^88^. Motif search was conducted by utilizing Fimo 5.4.1 of the MEME suite (https://meme-suite.org/meme/doc/fimo.html) against a random Markov background model which was created by running the *fasta-get-markov* command of the aforementioned suite, on random sequences that corresponded to the length and the chromosome of the query CTCF peaks, for each sample. Heatmaps were generated using deepTools^89^, while shared and condition-specific CTCF and SMC1A peaks were called using signal in the 100 bp around the summit of each peak (as calculated via SEACR).

### Micro-C and data analysis

Micro-C was performed using the Micro-C v1.0 kit in collaboration with Dovetail Genomics as per manufacturer’s instructions. Micro-C libraries (at least 3 per each biological replicate) that passed QC criteria were pooled and paired-end sequenced on a NovaSeq6000 platform (Illumina) to >600 million read pairs per replicate. Micro-C contact matrices were produced using Dovetail Genomics pipeline (https://micro-c.readthedocs.io/en/latest/fastq_to_bam.html). In brief, read pairs were mapped to human reference genome hg38 using BWA, after which low mapping quality (<40) reads and PCR duplicates were filtered out using the *MarkDuplicates* function in Picard tools (v2.20.7), and read coverage tracks (BigWig) were generated and normalized with the RPCG parameter using the *bamCoverage* function of deepTools2 v3.5.1^89^. Compartment boundaries for each sample corresponded to the 1 bp of adjacent bins on which compartment changed from A to B or from B to A. The interaction decay plot was created by using *cooltools* 0.5.1 (https://cooltools.readthedocs.io/en/latest/notebooks/contacts_vs_distance.html). The eigenvalues, needed for the saddle plots, were computed with the *cooltools call-compartments* command at 10-kbp resolution and the expected interactions were computed with cooltools compute-expected command at the same resolution. The saddle plot was created with cooltools compute-saddle using 100 digitized bins as described here: https://cooltools.readthedocs.io/en/latest/notebooks/compartments_and_saddles.html Finally, we used *coolpuppy* 0.9.5 (https://coolpuppy.readthedocs.io/en/latest/) to generate all aggregate plots. For loop calling, we used a multi-tool (HiCCUPS, SIP, and mustache) and a multi-resolution (5-and 10-kbp) approach as previously described^55,90^. Loop lists coming from each of the three different tools and across the two resolutions were merged using the *pgltools* intersect command with a distance tolerance of 1bp. This procedure results in considering loops that were called in adjacent bins across different resolutions or tools as being shared, while unique loops are considered those that exhibit a distance corresponding to at least one bin size (5-or 10-kbp) across the different loop-calling approaches. In cases of shared loops across the two resolutions, the 5kb resolution coordinates were kept for further analysis. In order to find condition-specific loops we furtherly annotated them with ICM-specific CTCF peaks. To detect ICM enriched CTCF peaks, we furtherly filtered peaks based on the control and ICM CUT&TAG signal enclosed in regions around the summits of ICM CTCF peaks. In more detail, we extracted the control and ICM depth-normalized CUT&TAG signal of regions 100 bp around the summits of peaks by utilizing the *multiBigwigSummary* command of Deeptools. The CTCF peaks that we considered in the downstream analysis were those that exhibited less than the mean control CUT&TAG signal with higher or equal to 1-fold difference compared to the corresponding ICM signal. 2628 ICM CTCF peaks fulfilled these criteria and were furtherly used to annotate both control and ICM loops. All intersections were performed using pgltools intersect1D without any distance tolerance for CTCF anchors. We considered loops as CTCF-associated when at least one of the anchors overlapped a CTCF peak of the subset described above. The rest of the loops were annotated as non-CTCF. We furtherly divided the loops into condition-specific and shared loops. Condition-specific loops had at least one unique anchor. This analysis was done, as described before, by utilizing the *pgltools* intersect command with 1 bp tolerance distance for both the shared and the unique loops. All code for the Micro-C analysis can be found at https://github.com/shuzhangcourage/Micro-C-CUT-tag/tree/v1.0.0.

### Single-cell RNA-seq

In brief proliferating, ICM-treated, and senescent IMR90s (8×10^5^ cells/condition) were grown to 80% confluency, harvested with trypsin and froze at-80°C. Single cell RNA-seq was performed using the 10X Genomics kit in collaboration with Active Motif. Libraries that passed QC criteria were paired-end sequenced to at least 250 million reads per library. All downstream analysis was performed using Seurat^91^.

### Statistical testing

*P*-values associated with Student’s *t*-tests, Fischer’s exact tests and with the Wilcoxon– Mann–Whitney tests were calculated using GraphPad (https://graphpad.com/). Unless otherwise stated, *P-* values < 0.01 were deemed as statistically significant.

### Data availability

All NGS data generated have been deposited to the NCBI Gene Expression Omnibus (GEO) as part of the GSE238256 SuperSeries (https://www.ncbi.nlm.nih.gov/geo/query/acc.cgi?acc=GSE238256). All imaging data is on the OSF (https://osf.io/n7qxc/?view_only=067396ac892542fc98767205f2063613).

---

**Supplementary Fig 1.**
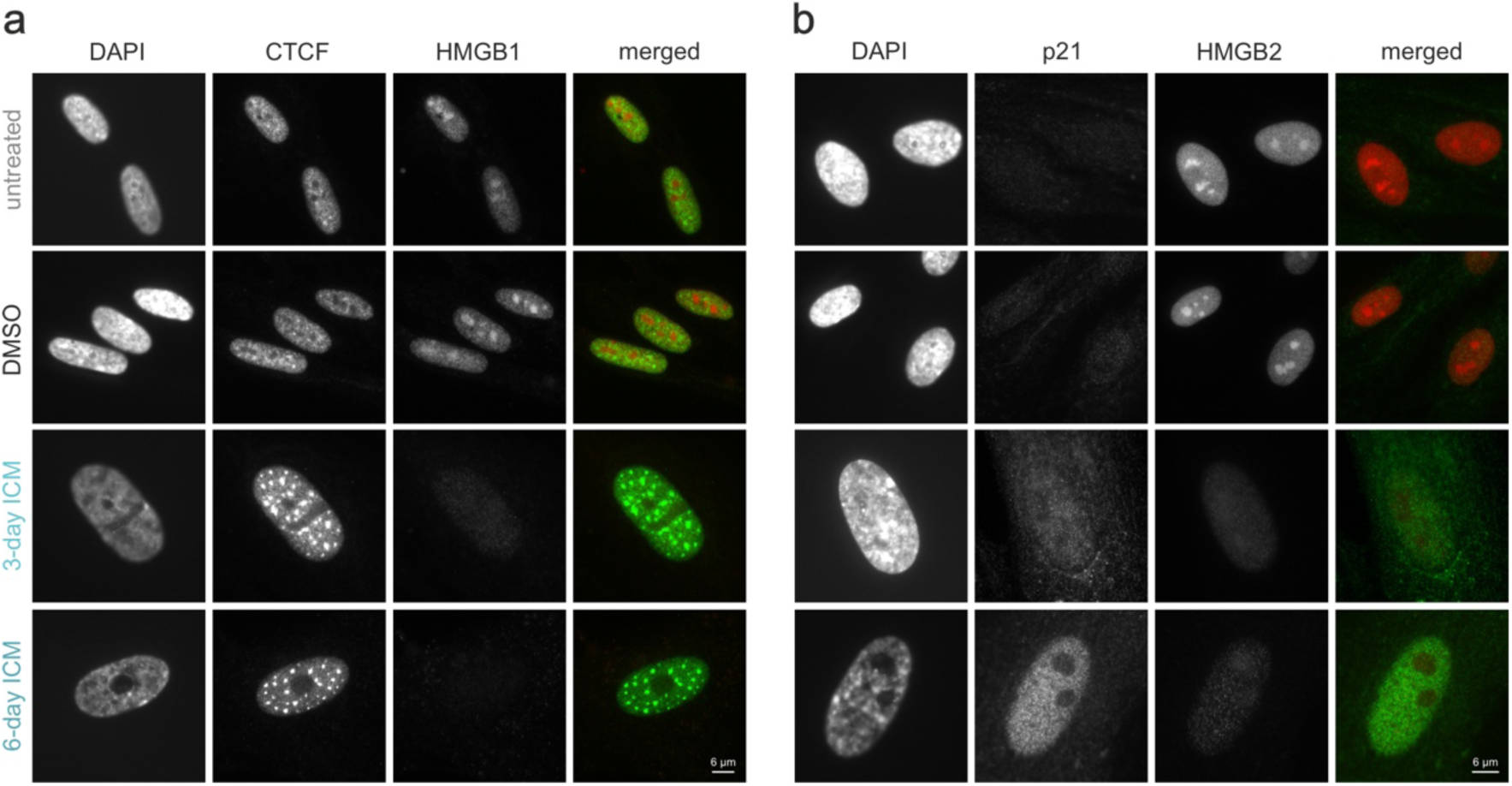
ICM treatment depletes HMGB1 and -B2 from IMR90 cell nuclei. **a.** Representative widefield images of proliferating, DMSO-, and 3-or 6-day ICM-treated IMR90 immunostained for CTCF and HMGB1, and counterstained with DAPI. Bar: 6 μm. **b.** As in panel A, but immunostained for p21 and HMGB2. Bar: 6 μm.

**Supplementary Fig 2.**
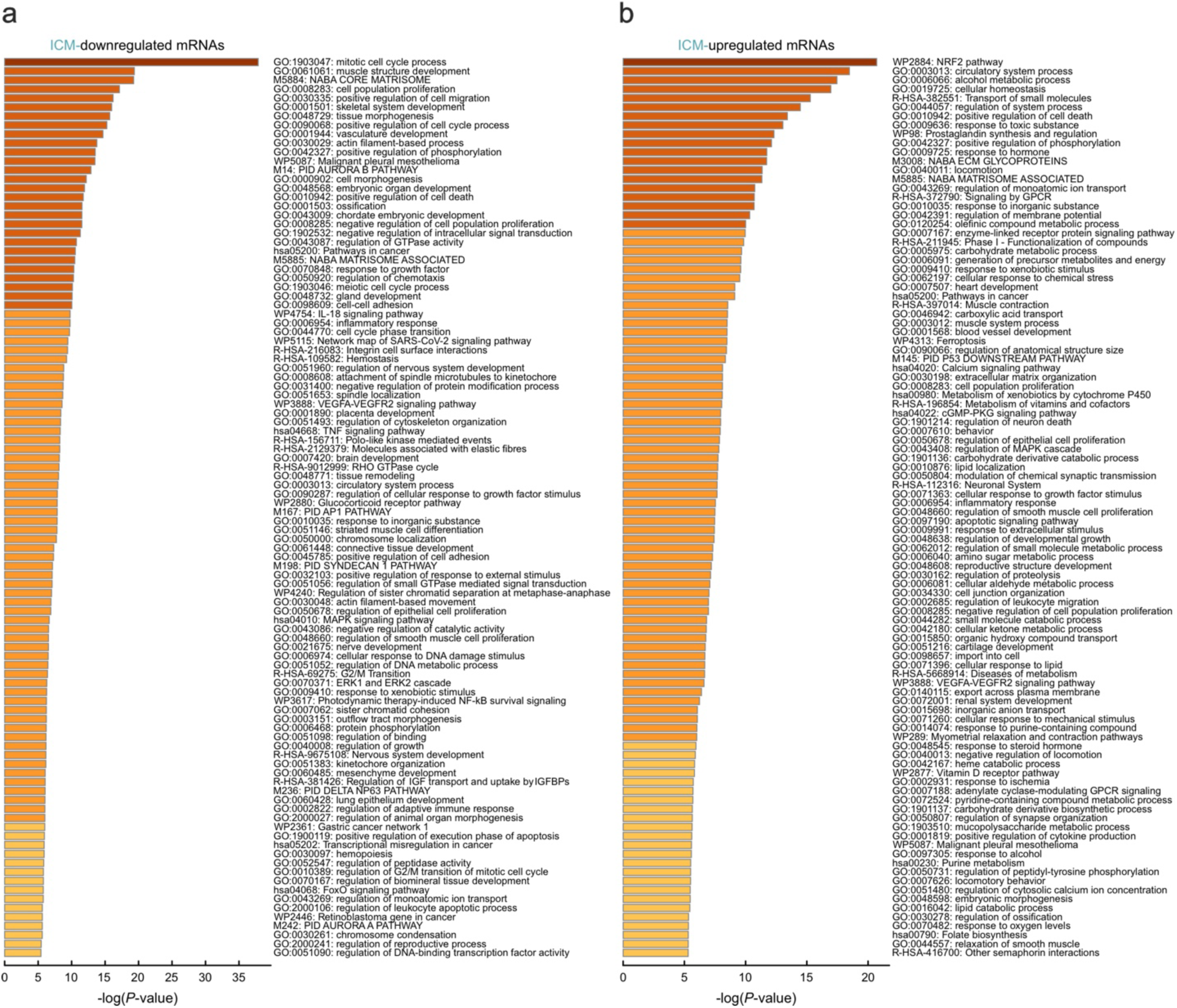
Effects of ICM treatment at the level of mRNA in IMR90. **a.** Top 100 GO terms/pathways associated with mRNAs downregulated following 6 days of 10 μΜ ICM treatment. **b.** As in panel A, but for ICM-upregulated mRNAs.

**Supplementary Fig 3.**
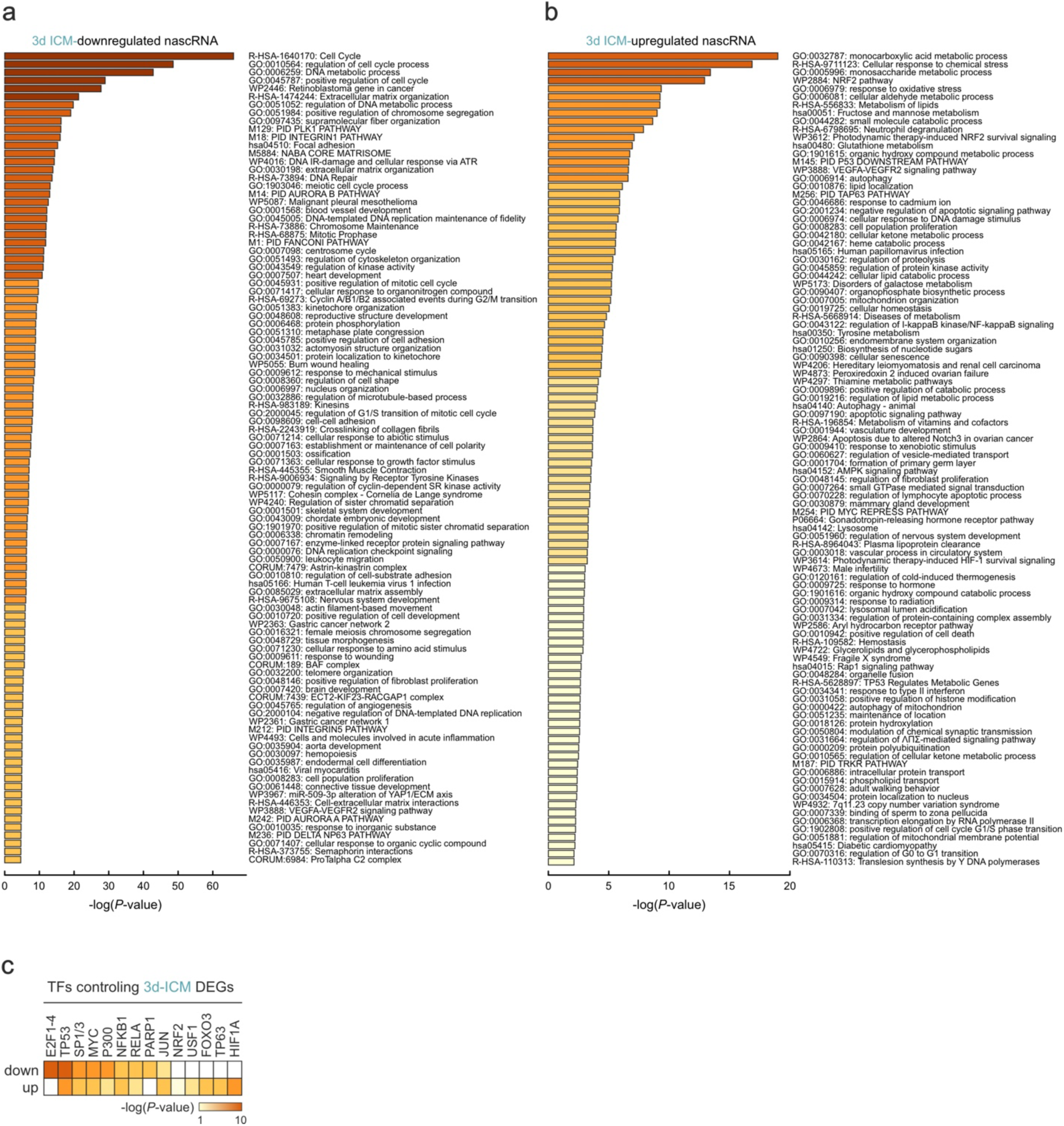
Effects of 3-day ICM treatment at the level of nascent RNA production in IMR90. **a.** Top 100 GO terms/pathways associated with nascent RNAs downregulated after 3 days of 10 μΜ ICM treatment. **b.** As in panel A, but for 3-day ICM-upregulated nascent RNAs. **c.** Heatmap showing transcription factors (TFs) predicted to bind ICM-regulated genes from panels A and B based on TTRUST motif enrichment.

**Supplementary Fig 4.**
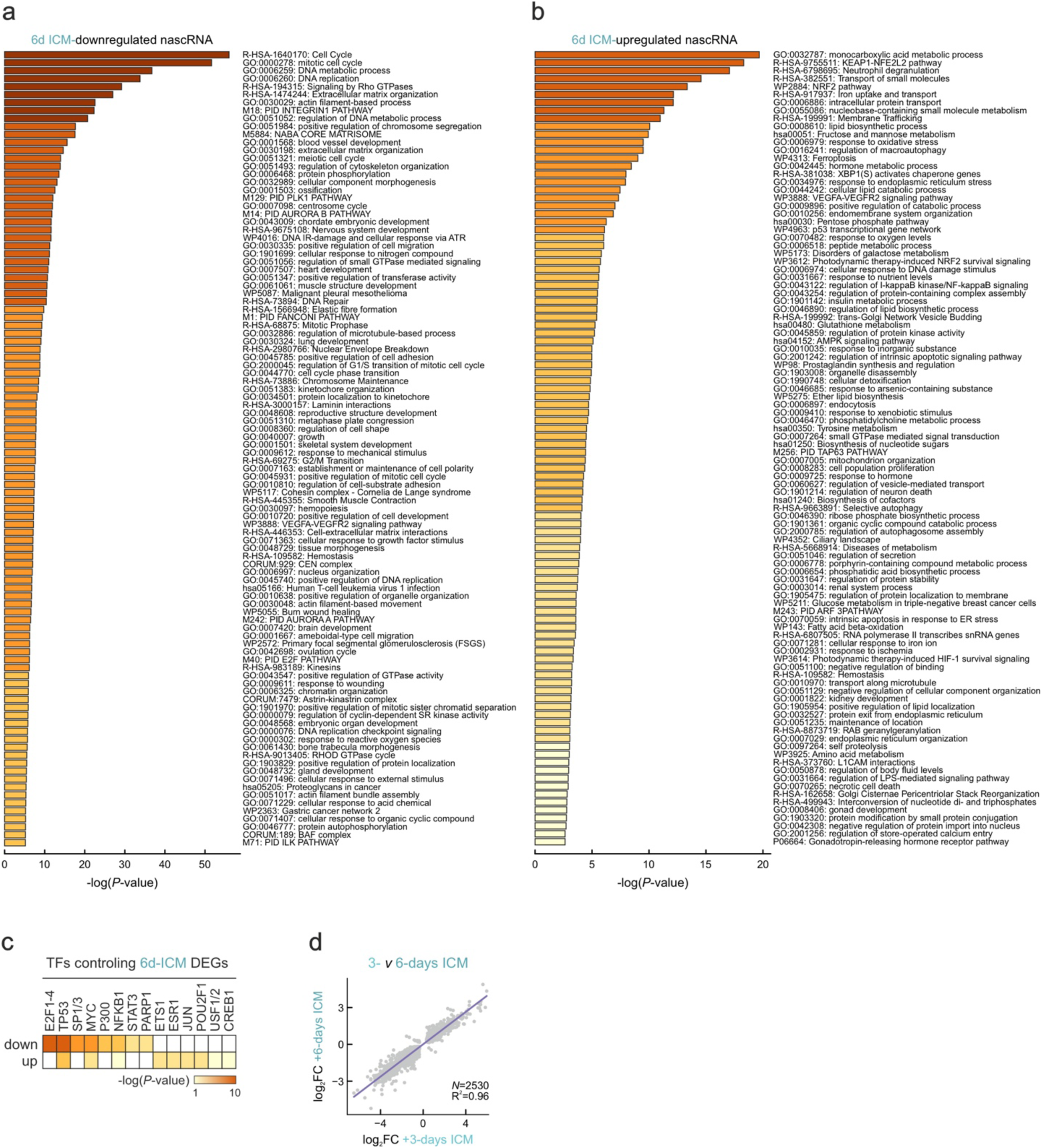
Effects of 6-dayICM treatment at the level of nascent RNA production in IMR90. **a.** Top 100 GO terms/pathways associated with nascent RNAs downregulated after 6 days of 10 μΜ ICM treatment. **b.** As in panel A, but for 6-day ICM-upregulated nascent RNAs. **c.** Heatmap showing transcription factors (TFs) predicted to bind ICM-regulated genes from panels A and B based on TTRUST motif enrichment. **d.** Plot correlating changes (log2FC) in nascent RNA levels of genes regulated after 3 and 6 days of ICM treatment. The number of the queried genes (*N*) and the deduced Spearman’s correlation coefficient (R2) are shown.

**Supplementary Fig 5.**
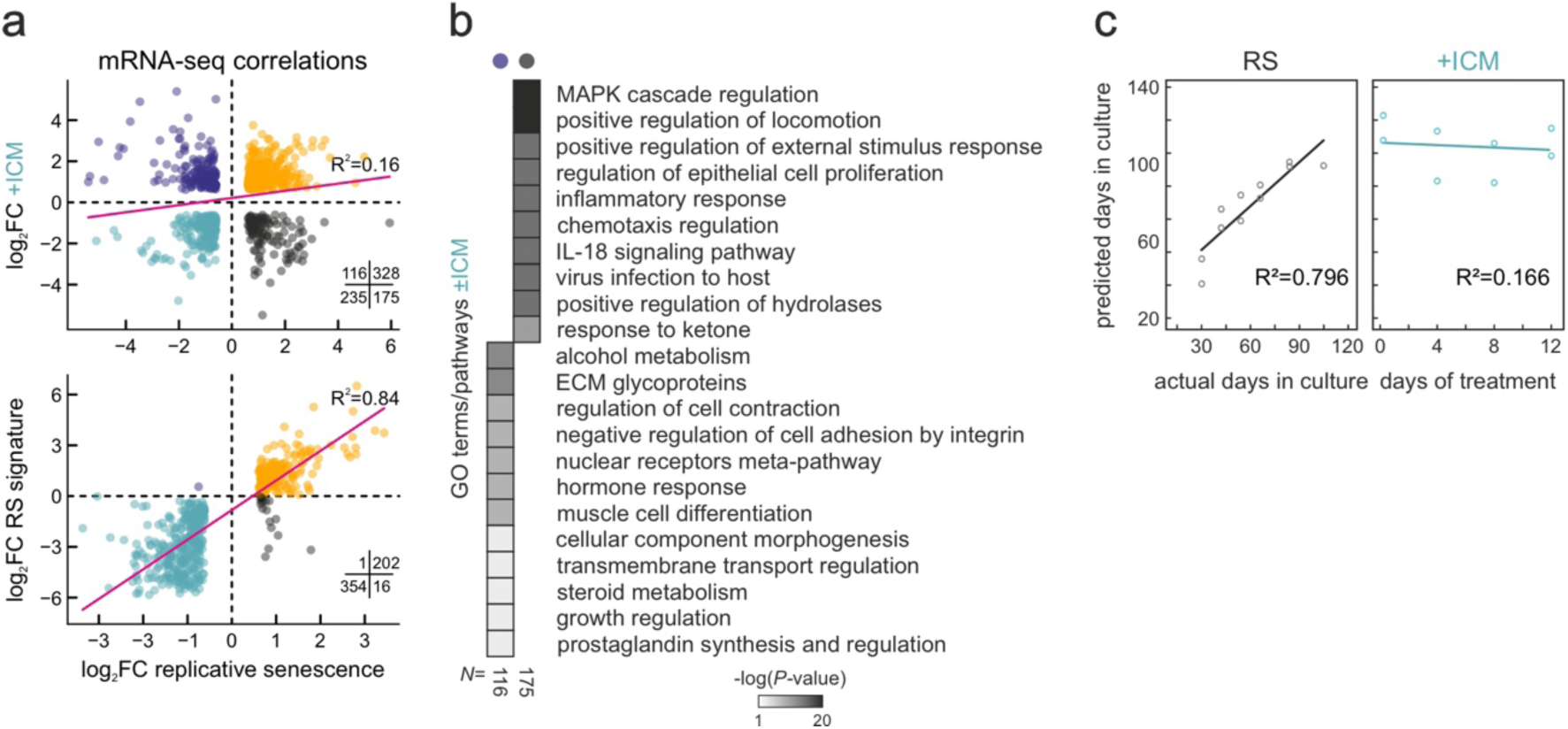
Transcriptional and methylation clock changes in 6-day ICM-treated IMR90. **a.** Top: Scatter plots showing correlation of differentially expressed mRNAs from ICM-treated and replicatively senescent IMR90. Bottom: As above, but for mRNAs from ICM-treated IMR90 and a consensus senescence signature. Spearman’s correlation coefficients (R2) and the number of genes in each quadrant (*N)* are shown. **b.** Heatmap of GO terms/pathways associated with the two diverging gene subsets from panel A (up in ICM and down in RS – purple; down in ICM and up in RS – black). The number of genes in each subset (*N*) is indicated. **c.** Plots correlating IMR90 passage predicted by methylation changes at six senescence-associated CpGs with actual passage (left) or days of ICM treatment (right). Spearman’s correlation coefficients (R2) are shown.

**Supplementary Fig 6.**
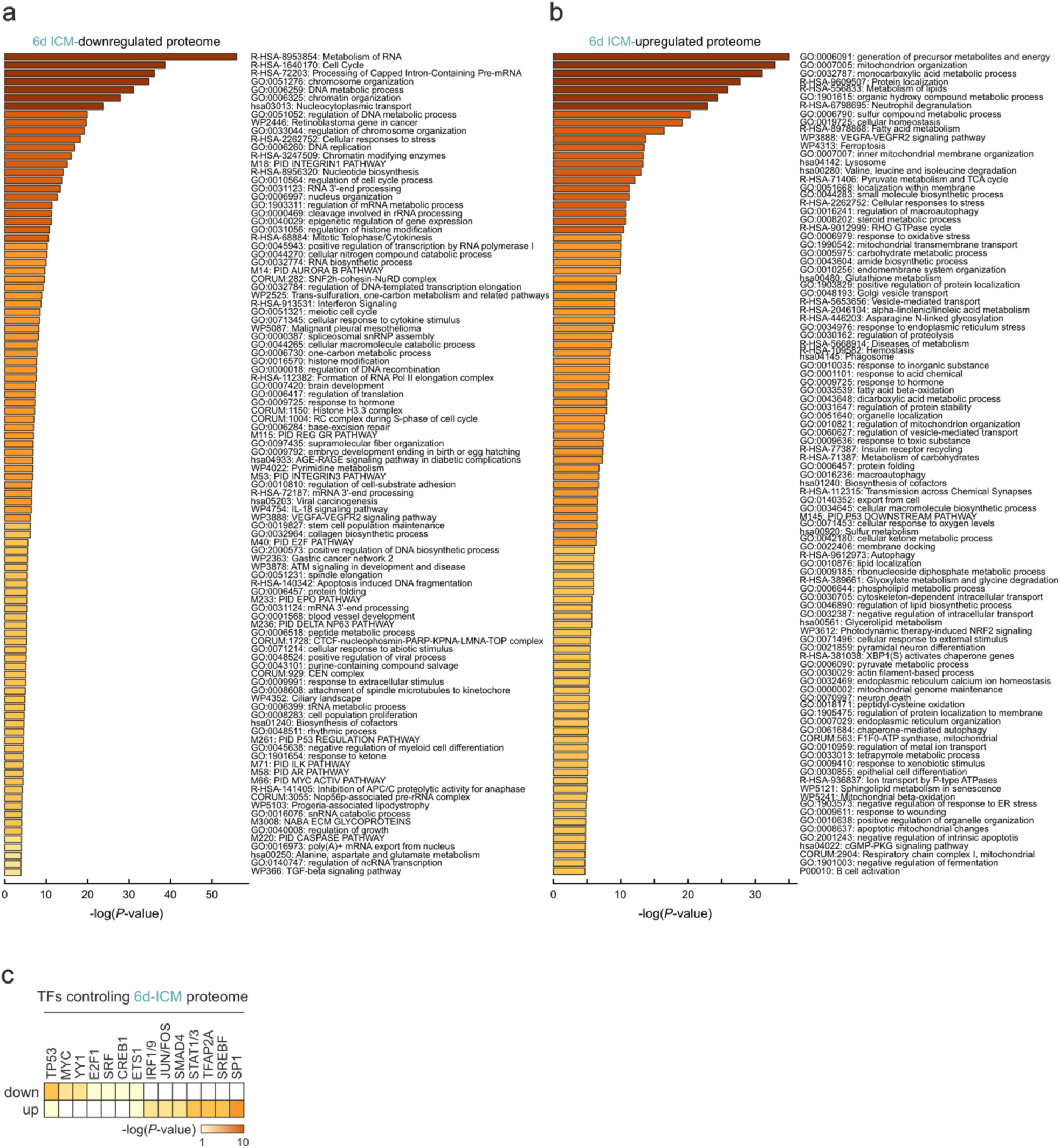
Effects of ICM treatment at the whole-cell proteome level in IMR90. **a.** Top 100 GO terms/pathways associated with proteins downregulated after 6 days of 10 μΜ ICM treatment. **b.** As in panel A, but for ICM-upregulated proteins. **c.** Heatmap showing transcription factors (TFs) predicted to bind the genes endoding the ICM-regulated genes from panels A and B based on TTRUST motif enrichment.

**Supplementary Table 1.**
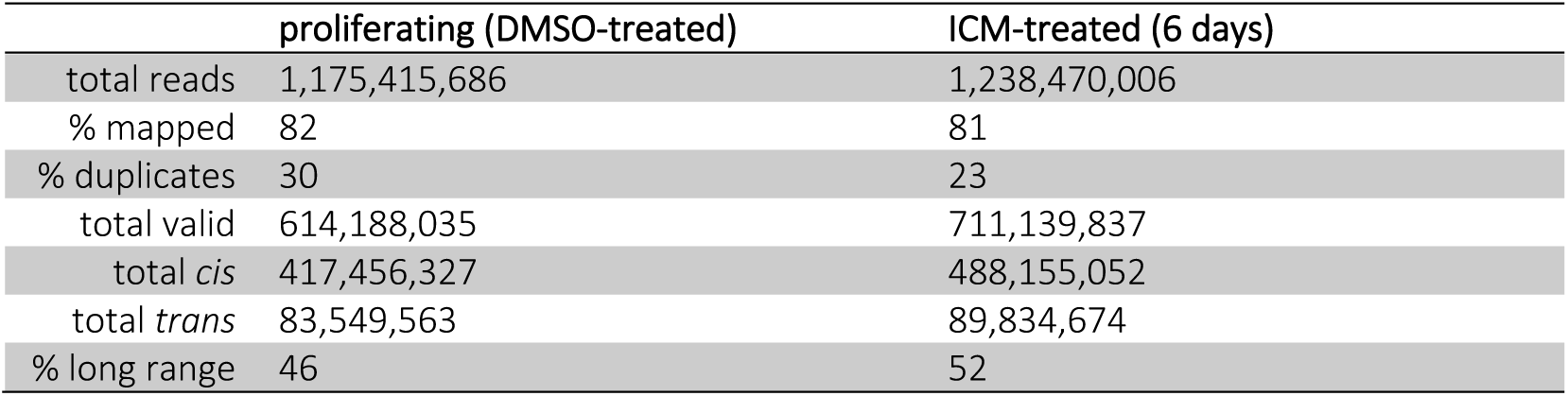
Basic mapping statistics of Micro-C experiments.

|  | proliferating (DMSO-treated) | ICM-treated (6 days) |
| --- | --- | --- |
| total reads | 1,175,415,686 | 1,238,470,006 |
| % mapped | 82 | 81 |
| % duplicates | 30 | 23 |
| total valid | 614,188,035 | 711,139,837 |
| total *cis* | 417,456,327 | 488,155,052 |
| total *trans* | 83,549,563 | 89,834,674 |
| % long range | 46 | 52 |

**Supplementary Table 2.**
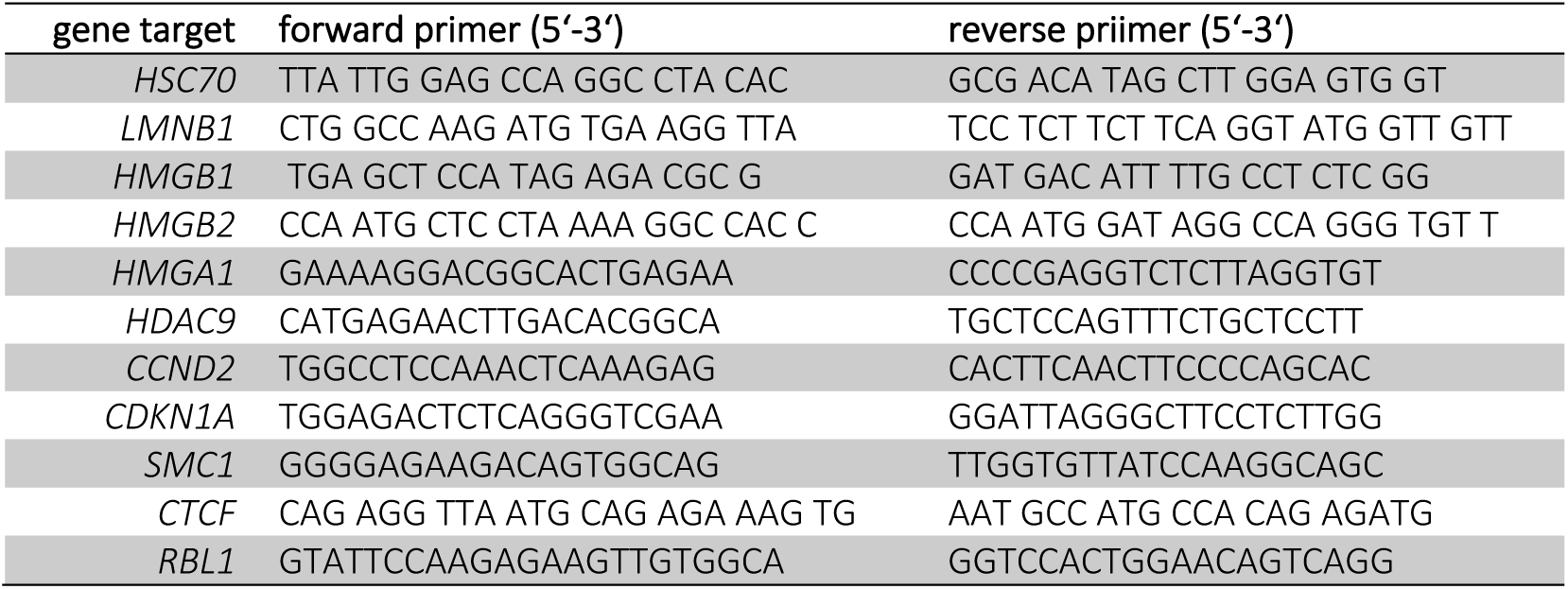
List of primers used in RT-qPCR experiments.

**Supplementary Table 3.**
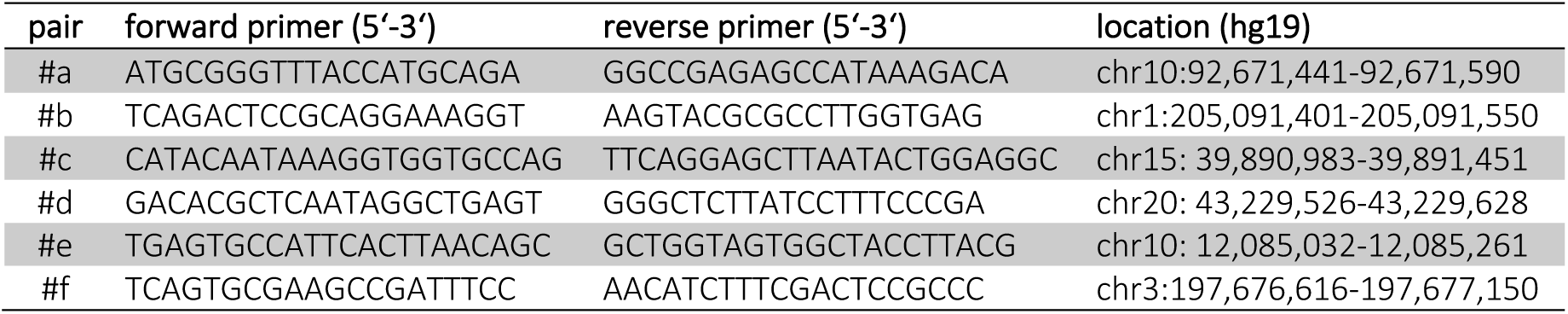
List of primers used in HMGB2 ChIP-qPCR experiments.

**Supplementary Table 4. Full list of peptides discovered by whole-cell proteomics and associated statistics. (Provided as an .*xlsx* file.)**

## References

1. López-Otín, C., Blasco, M. A., Partridge, L., Serrano, M. & Kroemer, G. The Hallmarks of Aging. Cell 153, 1194–1217 (2013).

2. Schmeer, C., Kretz, A., Wengerodt, D., Stojiljkovic, M. & Witte, O. W. Dissecting Aging and Senescence— Current Concepts and Open Lessons. Cells 2019, Vol. 8, Page 1446 8, 1446 (2019).

3. Gorgoulis, V. et al. Cellular Senescence: Defining a Path Forward. Cell 179, 813–827 (2019).

4. Baker, D. J. et al. Clearance of p16Ink4a-positive senescent cells delays ageing-associated disorders. Nat. 2011 4797372 479, 232–236 (2011).

5. Baker, D. J. et al. Naturally occurring p16Ink4a-positive cells shorten healthy lifespan. Nature 530, 184 (2016).

6. Wang, W. et al. A genome-wide CRISPR-based screen identifies KAT7 as a driver of cellular senescence. Sci. Transl. Med. 13, 2655 (2021).

7. Wang, L., Lankhorst, L. & Bernards, R. Exploiting senescence for the treatment of cancer. Nat. Rev. Cancer 2022 226 22, 340–355 (2022).

8. Campisi, J. Aging, Cellular Senescence, and Cancer. 10.1146/annurev-physiol-030212-183653 75, 685–705 (2013).

9. Hernandez-Segura, A. et al. Unmasking Transcriptional Heterogeneity in Senescent Cells. Curr. Biol. 27, 2652–2660.e4 (2017).

10. Narita, M. et al. Rb-mediated heterochromatin formation and silencing of E2F target genes during cellular senescence. Cell 113, 703–716 (2003).

11. Shah, P. P. et al. Lamin B1 depletion in senescent cells triggers large-scale changes in gene expression and the chromatin landscape. Genes Dev. 27, 1787–1799 (2013).

12. Sadaie, M. et al. Redistribution of the Lamin B1 genomic binding profile affects rearrangement of heterochromatic domains and SAHF formation during senescence. Genes Dev. 27, 1800–1808 (2013).

13. Boumendil, C., Hari, P., Olsen, K. C. F., Acosta, J. C. & Bickmore, W. A. Nuclear pore density controls heterochromatin reorganization during senescence. Genes Dev. 33, 144–149 (2019).

14. Sati, S. et al. 4D Genome Rewiring during Oncogene-Induced and Replicative Senescence. Mol. Cell 78, 522–538.e9 (2020).

15. Chandra, T. et al. Global reorganization of the nuclear landscape in senescent cells. Cell Rep. 10, 471–483 (2015).

16. Zhang, X. et al. The loss of heterochromatin is associated with multiscale three-dimensional genome reorganization and aberrant transcription during cellular senescence. Genome Res. 31, 1121–1135 (2021).

17. Cruickshanks, H. A. et al. Senescent cells harbour features of the cancer epigenome. Nat. Cell Biol. 2013 1512 15, 1495–1506 (2013).

18. Papantonis, A. HMGs as rheostats of chromosomal structure and cell proliferation. Trends Genet. 37, 986– 994 (2021).

19. Sofiadis, K. et al. HMGB1 coordinates SASP-related chromatin folding and RNA homeostasis on the path to senescence. Mol. Syst. Biol. 17, (2021).

20. Zirkel, A. et al. HMGB2 Loss upon Senescence Entry Disrupts Genomic Organization and Induces CTCF Clustering across Cell Types. Mol. Cell 70, 730–744.e6 (2018).

21. Criscione, S. W., Teo, Y. V. & Neretti, N. The Chromatin Landscape of Cellular Senescence. Trends Genet. 32, 751–761 (2016).

22. De Cecco, M. et al. Genomes of replicatively senescent cells undergo global epigenetic changes leading to gene silencing and activation of transposable elements. Aging Cell 12, 247–256 (2013).

23. Sen, P. et al. Spurious intragenic transcription is a feature of mammalian cellular senescence and tissue aging. Nat. aging 3, 402 (2023).

24. Debès, C. et al. Ageing-associated changes in transcriptional elongation influence longevity. Nat. 6167958 616, 814–821 (2023).

25. Olan, I. et al. Transcription-dependent cohesin repositioning rewires chromatin loops in cellular senescence. Nat. Commun. 2020 111 11, 1–14 (2020).

26. Acosta, J. C. et al. A complex secretory program orchestrated by the inflammasome controls paracrine senescence. Nat. Cell Biol. 2013 158 15, 978–990 (2013).

27. Laberge, R. M. et al. MTOR regulates the pro-tumorigenic senescence-associated secretory phenotype by promoting IL1A translation. Nat. Cell Biol. 2014 178 17, 1049–1061 (2015).

28. Kang, C. et al. The DNA damage response induces inflammation and senescence by inhibiting autophagy of GATA4. Science (80-.). 349, (2015).

29. Wiley, C. D. et al. Mitochondrial Dysfunction Induces Senescence with a Distinct Secretory Phenotype. Cell Metab. 23, 303–314 (2016).

30. Lopes-Paciencia, S. et al. The senescence-associated secretory phenotype and its regulation. Cytokine 117, 15–22 (2019).

31. Sun, Y., Coppé, J. P. & Lam, E. W. F. Cellular Senescence: The Sought or the Unwanted? Trends Mol. Med. 24, 871–885 (2018).

32. Sun, L., Yu, R. & Dang, W. Chromatin Architectural Changes during Cellular Senescence and Aging. Genes (Basel). 9, (2018).

33. Teo, Y. V. et al. Notch Signaling Mediates Secondary Senescence. Cell Rep. 27, 997–1007.e5 (2019).

34. Martin, L., Schumacher, L. & Chandra, T. Modelling the dynamics of senescence spread. Aging Cell e13892 (2023) doi:10.1111/ACEL.13892.

35. Wiley, C. D. et al. Analysis of individual cells identifies cell-to-cell variability following induction of cellular senescence. Aging Cell 16, 1043–1050 (2017).

36. Chan, M. et al. Novel Insights from a Multiomics Dissection of the Hayflick Limit. Elife 11, (2022).

37. Lee, S. et al. A small molecule binding HMGB1 and HMGB2 inhibits microglia-mediated neuroinflammation. Nat. Chem. Biol. 10, 1055–1060 (2014).

38. Hayflick, L. & Moorhead, P. S. The serial cultivation of human diploid cell strains. Exp. Cell Res. 25, 585–621 (1961).

39. Harley, C. B., Futcher, A. B. & Greider, C. W. Telomeres shorten during ageing of human fibroblasts. Nature 345, 458–460 (1990).

40. Krtolica, A., Parrinello, S., Lockett, S., Desprez, P. Y. & Campisi, J. Senescent fibroblasts promote epithelial cell growth and tumorigenesis: a link between cancer and aging. Proc. Natl. Acad. Sci. U. S. A. 98, 12072– 12077 (2001).

41. Coppe, J. P., Kauser, K., Campisi, J. & Beauséjour, C. M. Secretion of Vascular Endothelial Growth Factor by Primary Human Fibroblasts at Senescence. J. Biol. Chem. 281, 29568–29574 (2006).

42. Demaria, M. et al. An Essential Role for Senescent Cells in Optimal Wound Healing through Secretion of PDGF-AA. Dev. Cell 31, 722–733 (2014).

43. Dimri, G. P. et al. A biomarker that identifies senescent human cells in culture and in aging skin in vivo. Proc. Natl. Acad. Sci. U. S. A. 92, 9363–9367 (1995).

44. Coppé, J. P., Desprez, P. Y., Krtolica, A. & Campisi, J. The senescence-associated secretory phenotype: the dark side of tumor suppression. Annu. Rev. Pathol. 5, 99–118 (2010).

45. Han, H. et al. TRRUST v2: an expanded reference database of human and mouse transcriptional regulatory interactions. Nucleic Acids Res. 46, D380–D386 (2018).

46. Narita, M. et al. A Novel Role for High-Mobility Group A Proteins in Cellular Senescence and Heterochromatin Formation. Cell 126, 503–514 (2006).

47. Parry, A. J. et al. NOTCH-mediated non-cell autonomous regulation of chromatin structure during senescence. Nat. Commun. 2018 91 9, 1–15 (2018).

48. Doubleday, P. F., Fornelli, L. & Kelleher, N. L. Elucidating Proteoform Dynamics Underlying the Senescence Associated Secretory Phenotype. J. Proteome Res. 19, 938–948 (2020).

49. Caudron-Herger, M., Cook, P. R., Rippe, K. & Papantonis, A. Dissecting the nascent human transcriptome by analysing the RNA content of transcription factories. Nucleic Acids Res. 43, e95 (2015).

50. Rai, T. S. et al. HIRA orchestrates a dynamic chromatin landscape in senescence and is required for suppression of neoplasia. Genes Dev. 28, 2712–2725 (2014).

51. Franzen, J. et al. Senescence-associated DNA methylation is stochastically acquired in subpopulations of mesenchymal stem cells. Aging Cell 16, 183–191 (2017).

52. Papaspyropoulos, A., et al. Decoding of translation-regulating entities reveals heterogeneous translation deficiency patterns in cellular senescence. Aging Cell In Press, (2023).

53. Davalos, A. R. et al. p53-dependent release of Alarmin HMGB1 is a central mediator of senescent phenotypes. J. Cell Biol. 201, 613–629 (2013).

54. Mizi, A., Zhang, S. & Papantonis, A. Genome folding and refolding in differentiation and cellular senescence. Curr. Opin. Cell Biol. 67, 56–63 (2020).

55. Hsieh, T.-H. S. et al. Resolving the 3D Landscape of Transcription-Linked Mammalian Chromatin Folding. (2020) doi:10.1016/j.molcel.2020.03.002.

56. Criscione, S. W. et al. Reorganization of chromosome architecture in replicative cellular senescence. Sci. Adv. 2, (2016).

57. Hansen, A. S., Pustova, I., Cattoglio, C., Tjian, R. & Darzacq, X. CTCF and cohesin regulate chromatin loop stability with distinct dynamics. Elife 6, (2017).

58. Zhang, S., Übelmesser, N., Barbieri, M. & Papantonis, A. Enhancer–promoter contact formation requires RNAPII and antagonizes loop extrusion. Nat. Genet. 2023 555 55, 832–840 (2023).

59. Kirschner, K., Rattanavirotkul, N., Quince, M. F. & Chandra, T. Functional heterogeneity in senescence. Biochem. Soc. Trans. 48, 765–773 (2020).

60. Robbins, P. D. et al. Annual Review of Pharmacology and Toxicology Senolytic Drugs: Reducing Senescent Cell Viability to Extend Health Span. (2020) doi:10.1146/annurev-pharmtox-050120.

61. Chaib, S., Tchkonia, T. & Kirkland, J. L. Cellular senescence and senolytics: the path to the clinic. Nat. Med. 2022 288 28, 1556–1568 (2022).

62. Galiana, I. et al. Preclinical antitumor efficacy of senescence-inducing chemotherapy combined with a nanoSenolytic. J. Control. Release 323, 624–634 (2020).

63. Meang, M. K., Kim, S., Kim, I. H., Kim, H. S. & Youn, B. S. A Small Molecule That Promotes Cellular Senescence Prevents Fibrogenesis and Tumorigenesis. Int. J. Mol. Sci. 23, (2022).

64. Browder, K. C. et al. In vivo partial reprogramming alters age-associated molecular changes during physiological aging in mice. Nat. Aging 2, 243–253 (2022).

65. Roux, A. E. et al. The complete cell atlas of an aging multicellular organism. bioRxiv 2022.06.15.496201 (2022) doi:10.1101/2022.06.15.496201.

66. Salminen, A., Ojala, J., Kaarniranta, K. & Kauppinen, A. Mitochondrial dysfunction and oxidative stress activate inflammasomes: impact on the aging process and age-related diseases. Cell. Mol. Life Sci. 2012 6918 69, 2999–3013 (2012).

67. Vénéreau, E., Ceriotti, C. & Bianchi, M. E. DAMPs from cell death to new life. Front. Immunol. 6, 159317 (2015).

68. Chung, H. et al. High mobility group box 1 secretion blockade results in the reduction of early pancreatic islet graft loss. Biochem. Biophys. Res. Commun. 514, 1081–1086 (2019).

69. Kim, Y. H. et al. Inflachromene inhibits autophagy through modulation of Beclin 1 activity. J. Cell Sci. 131, (2018).

## Supplementary references

70. Bucevičius, J., Kostiuk, G., Gerasimaitė, R., Gilat, T. & Lukinavičius, G. Enhancing the biocompatibility of rhodamine fluorescent probes by a neighbouring group effect. Chem. Sci. 11, 7313–7323 (2020).

71. Schindelin, J. et al. Fiji: an open-source platform for biological-image analysis. *Nat. Methods* 2012 97 9, 676–682 (2012).

72. Lukinavičius, G. et al. SiR–Hoechst is a far-red DNA stain for live-cell nanoscopy. Nat. Commun. 6, 1–7 (2015).

73. Harris, C. R. et al. Array programming with NumPy. Nat. 2020 5857825 585, 357–362 (2020).

74. Van Der Walt, S. et al. Scikit-image: Image processing in python. PeerJ 2014, e453 (2014).

75. Pedregosa, F. et al. Scikit-learn: Machine Learning in Python. J. Mach. Learn. Res. 12, 2825–2830 (2011).

76. Hörl, D. et al. BigStitcher: reconstructing high-resolution image datasets of cleared and expanded samples. Nat. Methods 2019 169 16, 870–874 (2019).

77. Heckenbach, I. et al. Nuclear morphology is a deep learning biomarker of cellular senescence. Nat. Aging 2022 28 2, 742–755 (2022).

78. Dobin, A. et al. STAR: ultrafast universal RNA-seq aligner. Bioinformatics 29, 15–21 (2013).

79. Liao, Y., Smyth, G. K. & Shi, W. featureCounts: an efficient general purpose program for assigning sequence reads to genomic features. Bioinformatics 30, 923–930 (2014).

80. Risso, D., Ngai, J., Speed, T. P. & Dudoit, S. Normalization of RNA-seq data using factor analysis of control genes or samples. Nat. Biotechnol. 32, 896–902 (2014).

81. Love, M. I., Huber, W. & Anders, S. Moderated estimation of fold change and dispersion for RNA-seq data with DESeq2. Genome Biol. 15, (2014).

82. Zhou, Y. et al. Metascape provides a biologist-oriented resource for the analysis of systems-level datasets. Nat. Commun. 10, (2019).

83. Melnik, S. et al. Isolation of the protein and RNA content of active sites of transcription from mammalian cells. Nat. Protoc. 2016 113 11, 553–565 (2016).

84. Madsen, J. G. S. et al. iRNA-seq: computational method for genome-wide assessment of acute transcriptional regulation from total RNA-seq data. Nucleic Acids Res. 43, e40 (2015).

85. Ivanov, I. P. et al. Polyamine Control of Translation Elongation Regulates Start Site Selection on Antizyme Inhibitor mRNA via Ribosome Queuing. Mol. Cell 70, 254–264.e6 (2018).

86. Oertlin, C. et al. Generally applicable transcriptome-wide analysis of translation using anota2seq. Nucleic Acids Res. 47, E70 (2019).

87. Langmead, B. & Salzberg, S. L. Fast gapped-read alignment with Bowtie 2. Nat. Methods 2012 94 9, 357– 359 (2012).

88. Grant, C. E., Bailey, T. L. & Noble, W. S. FIMO: scanning for occurrences of a given motif. Bioinformatics 27, 1017–1018 (2011).

89. Ramírez, F., Dündar, F., Diehl, S., Grüning, B. A. & Manke, T. deepTools: a flexible platform for exploring deep-sequencing data. Nucleic Acids Res. 42, (2014).

90. Krietenstein, N. et al. Ultrastructural details of mammalian chromosome architecture. Mol. Cell 78, 554 (2020).

91. Satija, R., Farrell, J. A., Gennert, D., Schier, A. F., & Regev, A. (2015). Spatial reconstruction of single-cell gene expression data. Nat. Biotech. 33, 495–502 (2015)

